# Acute adenoviral infection elicits an arrhythmogenic substrate prior to myocarditis

**DOI:** 10.1101/2022.11.29.518421

**Authors:** Rachel L. Padget, Michael J. Zeitz, Grace A. Blair, Xiaobo Wu, Michael D. North, Mira T. Tanenbaum, Kari E. Stanley, Chelsea M. Phillips, D. Ryan King, Samy Lamouille, Robert G. Gourdie, Gregory S. Hoeker, Sharon A. Swanger, Steven Poelzing, James W. Smyth

## Abstract

**Background:** Viral cardiac infection represents a significant clinical challenge encompassing several etiological agents, disease stages, complex presentation, and a resulting lack of mechanistic understanding. Myocarditis is a major cause of sudden cardiac death in young adults, where current knowledge in the field is dominated by later disease phases, and pathological immune responses. However, little is known regarding how infection can acutely induce an arrhythmogenic substrate prior to significant immune responses. Adenovirus is a leading cause of myocarditis, but due to species-specificity, models of infection are lacking and it is not understood how adenoviral infection may underlie sudden cardiac arrest. Mouse Adenovirus Type-3 (MAdV-3) was previously reported as cardiotropic, yet has not been utilized to understand mechanisms of cardiac infection and pathology.

**Methods:** We have developed MAdV-3 infection as a model to investigate acute cardiac infection and molecular alterations to the infected heart prior to an appreciable immune response or gross cardiomyopathy.

**Results:** Optical mapping of infected hearts exposes decreases in conduction velocity concomitant with increased Cx43Ser368 phosphorylation, a residue known to regulate gap junction function. Hearts from animals harboring a phospho-null mutation at Cx43Ser368 are protected against MAdV-3 induced conduction velocity slowing. Additional to gap junction alterations, patch clamping of MAdV-3-infected adult mouse ventricular cardiomyocytes reveals prolonged action potential duration as a result of decreased *I*_K1_ and *I*_Ks_ current density. Turning to human systems, we find human adenovirus type-5 (HAdV-5) increases phosphorylation of Cx43Ser368 and disrupts synchrony in human induced pluripotent stem cell-derived cardiomyocytes (iPSC-CMs), indicating common mechanisms with our mouse whole heart and adult cardiomyocyte data.

**Conclusions:** Together, these findings demonstrate that adenoviral infection creates an arrhythmogenic substrate through direct targeting of gap junction and ion channel function in the heart. Such alterations are known to precipitate arrhythmias and likely contribute to sudden cardiac death in acutely infected patients.

## Introduction

Viral infection of the heart results in significant morbidity and mortality, representing a major cause of sudden cardiac death (SCD) in young adults where up to a 42% causative incidence of viral-associated myocarditis is reported^1–3^. Viral myocarditis is classified into distinct phases of disease, encompassing an acute phase of active infection where direct viral-mediated damage to myocardial cells occurs and may subsequently progress to a chronic disease phase involving the host immune response^4^. Chronic viral myocarditis can then advance to dilated cardiomyopathy (DCM) and heart failure, with limited clinical options available bar the supportive care and potential need for transplant shared with other forms of end-stage cardiac disease^5,6^. Clinical myocarditis data are understandably dominated by chronic disease, where host immune responses to the initial infection are understood to precipitate pathological remodeling to HF^7^. The molecular mechanisms of acute cardiac infection, and specifically how active viral replication in the myocardium precipitates arrhythmia prior to an appreciable immune response, are not well understood. Indeed, individuals with such direct active cardiac involvement often do not survive the sudden cardiac arrest event triggered by infection and/or do not present with altered cardiac remodeling and function by echocardiography, for example^8,9^. A better understanding of how acute viral infection alters cardiac electrophysiology and function prior to immune infiltration is key in the development of interventions against arrhythmogenesis, in addition to identifying pathways underlying progression to chronic viral myocarditis for therapeutic targeting and/or early diagnosis.

Arrhythmias, while rare, are the leading cause of sudden cardiac arrest in children and young adults^3,10,11^. Hypertrophic obstructive cardiomyopathy and congenital abnormalities are significant contributors to SCD in such cases, but here we investigate how acute viral infection can elicit pathological remodeling preceding SCD in hearts otherwise devoid of such apparent cardiomyopathy. Normal cardiac electrophysiological function is achieved primarily through two major components: ion channels localized to specific nanodomains of the sarcolemma, and cell-to-cell electrical coupling at the intercalated disc (ID)^12–14^. The proteins comprising the major cardiac ion channels responsible for generation of the action potential (AP), including Ca_V_1.2, Na_V_1.5, Kir2.1, K_V_7.1, and K_V_11.1, are subject to dynamic regulation affecting their localization and function. Pathological alterations to the AP, such as prolonged duration due to perturbed channel function, can lead to lethal arrythmias^15^. IDs encompass fascia adherens, gap junctions, the perinexus, desmosomes, and an enrichment of several ion channels to effect mechanical and electrical coupling between adjacent cardiomyocytes^16–21^. Subcellular remodeling of such ID-resident structures and channels is well described as underlying arrhythmogenesis across a breadth of cardiac disease states, including viral infection^22–27^.

A rapid and modifiable mechanism of regulating structures at the ID is dynamic phosphorylation of target proteins. A common outcome of increased phosphorylation of gap junctions and ion channels is a positive or negative effect on function^28–30^. In addition to direct impacts, indirect consequences include regulation of ion channel density. For example, increased phosphorylation by PKC on KCNE1 can impair trafficking of the KCNQ1/KCNE1 complex forming the ion channel, K_V_7.1, and therefore reduce *I*_Ks_^31,32^. Changes to gap junction intracellular communication (GJIC) are established in underlying arrhythmias and SCD^33^.

Twenty-one human connexins have been identified, with connexin43 (Cx43) being the most ubiquitously expressed throughout the body while also being the primary connexin of the ventricular working myocardium^34^. In addition to electrical coupling, gap junctions play a critical role in propagation of innate and adaptive antiviral immune responses, providing justification for pathogens to target such structures^35–37^. Indeed, we have previously reported Cx43 is specifically down regulated at the transcriptional level and functionally targeted through altered phosphorylation during adenoviral infection in human epithelial cells and induced pluripotent stem cell-derived cardiomyocytes (HiPSC-CMs)^38^.

The most common viral families identified as causative agents in SCD and development of viral myocarditis include adenoviruses, enteroviruses (Coxsackievirus B3, CVB3), herpesviruses, and parvoviruses, with enteroviruses representing the most intensely researched^10,39,40^. SARS-CoV-2 infection has also been implicated both directly and indirectly in myocarditis development^40,41^. Regarding enteroviruses, CVB3 is a non-linear, positive-sense single-stranded RNA (ssRNA) virus from the picornavirus family commonly associated with childhood illness and a significant causative agent of viral myocarditis in all ages^42^. The ability of CVB3 to viably infect and replicate across species has resulted in its development as a valuable, and arguably leading, research model for viral myocarditis^27,43,44^. Indeed, the majority of viral myocarditis research has been conducted using CVB3 with an emphasis on the chronic phase of disease^7^. Other viruses known to cause myocarditis, such as adenoviruses, have starkly different biology to CVB3, perhaps most obviously is their double stranded DNA (dsDNA) genomes requiring the virus to colonize the cellular nucleus and uncouple cell cycle regulation for a significantly longer life cycle than a positive-sense RNA virus^42,45,46^. Such differences likely elicit virus-specific pathologies during the acute stage of viral myocarditis, illustrating the need to investigate virus family-specific disease progress and pathology beyond CVB3.

Adenoviruses are non-enveloped, dsDNA viruses, and include over 50 serotypes (now ‘types’) capable of human infection where they cause a range of illnesses from mild respiratory disease to more severe and potentially fatal conditions such as myocarditis^47^. Tissue tropism is classically attributed to type-specific host cell receptor binding and/or permissiveness of specific cell types to adenoviral replication^48^. Human adenovirus types 2 and 5 (HAdV-2 and −5) are the most predominant causes of adenoviral myocarditis, although additional types are increasingly reported^49–51^. Early adenoviral proteins mimic growth factor signaling and induce cell cycle entry to take advantage of DNA synthesis machinery and replicate viral genomes^46,52,53^. These same signaling pathways target GJIC, as we have previously demonstrated, and likely also impact the cardiac ion channels that shape the AP^28,31,38,54^. Unlike CVB3 however, many HAdVs cannot replicate effectively in mice, resulting in a dearth of understanding regarding adenoviral cardiotropism and virus-specific effects on the heart due to this stringent adenoviral species-specificity^47,55^. Mouse adenovirus type 1 (MAdV-1) has historically been employed in modeling respiratory disease and encephalitis, and while not a ‘cardiotropic’ virus, additional studies have successfully modeled adenoviral myocarditis in mice using MAdV-1, with an emphasis on inflammation^56–59^. In 2008, mouse adenovirus type-3 (MAdV-3) was isolated as distinct from the previously described MAdV-1 and MAdV-2, and reported to be cardiotropic^60,61^. Development of MAdV-3 as a model for myocarditis, or use of this virus in investigation mechanisms of arrhythmogenesis during acute cardiac infection has been limited, however.

In this study, we successfully employ MAdV-3 to model acute viral cardiac infection in adult mice and confirm similar targeting of Cx43 to that which we previously observed with human adenovirus^38^. While we do report induction of a cardiac antiviral immune response at the RNA level and increases in infiltrating macrophages, we find slowed conduction velocity in infected hearts and prolonged AP duration (APD) in infected cardiomyocytes; two pro-arrhythmogenic phenotypes occurring prior to appreciable adaptive immune cell infiltration or development of any gross cardiomyopathy. Importantly, our cell-based studies reveal electrophysiological and gap junction disturbances independent of a host immune system. Turning to studies in isolated primary adult cardiomyocytes we identify perturbation of *I*_KS_ and *I*_K1_ underlies prolonged APD during adenoviral infection. MAdV-3 infection therefore represents a viable and relevant model of acute myocarditis, and targets cardiac conduction through the same mechanisms we have previously found in cell-based human adenovirus studies. While we focus on an acute stage of 7 days post infection where presumably some inflammatory processes and cytokine effects will have initiated, significant adaptive immune cell infiltration and/or cardiomyocyte damage is not occurring at a detectable level. Our data demonstrate that acutely infected hearts harbor dangerous electrophysiological alterations at the molecular level and reveal, for the first time, how adenoviral infection can precipitate such pathological subcellular remodeling prior to cardiomyopathy and development of inflammatory myocarditis.

## Materials and Methods

All materials and methods are described in the Supplemental Material and Major Resources Table.

## Results

### Mouse adenovirus type-3 is cardiotropic and does not invoke an appreciable infiltrative cardiac immune response during acute infection

We previously reported specific targeting of Cx43 expression and gap junction function during adenoviral infection^38^. Such alterations in Cx43 would significantly impact cardiac electrophysiology directly through affecting GJIC, while potentially indirectly impacting sodium channels and/or other key components of arrhythmogenesis^19,21,62,63^. Despite significant incidence of adenoviral myocarditis, establishment of an adult mouse disease model has been elusive due to species-specificity^55^. We first sought to examine the potential of MAdV-3 as a model of acute cardiac infection, given the prior reports of cardiotropism^60^. To test this, 8-12 week old male and female C57BL/6 mice were inoculated with 5 × 10^5^ i.u. MAdV-3 via retro-orbital injection. Animals were sacrificed and organs harvested 7 d.p.i. for histology and DNA isolation. Using qPCR we find MAdV-3 viral genomes significantly enriched only within cardiac tissue and spleen (log fold-change heart relative to lung 2.1, spleen 0.78), confirming cardiotropism (Figure 1A). A similar time-course over 7 days focusing on cardiac tissue alone reveals levels of viral genomes plateauing from 3-7 d.p.i., consistent with active viral replication in the heart during this time (Figure 1B). No weight loss or apparent evidence of malaise (e.g. hunching, fur-ruffling) were observed in infected animals during the 7 day time course (data not shown). We then employed RNA sequencing of control and MadV3 infected ventricular tissue from mice to assess global transcriptional changes (Figure 1C). MadV3 infected ventricles had 321 significant differentially expressed genes in comparison to controls. Nearly all (99%) differentially expressed genes were upregulated in response to viral infection. Differentially expressed genes were hierarchically clustered using the clustergrammer^64^ web program and are presented as a heatmap. To determine enriched gene ontology terms, over-representation analysis was performed with the significant differentially expressed genes using ClusterProfiler^65^. The top 10 most significant gene ontology terms for biological processes are shown sorted by gene ratio. Gene ontology of biological processes indicated an enrichment of immune related genes comprising both innate and adaptive immune responses in MAdV-3 infected ventricular tissue. Histopathology, however, revealed no gross pathology or appreciable infiltration of leukocytes in infected cardiac tissue (Figure S1A). To measure infiltration more specifically, we performed confocal microscopy immunofluorescence labeling of ventricular myocardial cryosections for the major immune cell markers: F4/80 (macrophages), CD3 (T-cells), and CD45R/B220 (B-cells, NK cells, CTLs). While increased numbers of F4/80+ cells were detected in MAdV-3 infected hearts, no changes to CD3+ or CD45R+ cell numbers were found in ventricular tissues, with CD3+ cells being notably rare (Figure 1D,E; Figure S1B,C). A lack of direct cardiomyocyte damage and presence of a cardiac inflammatory response were confirmed by serum cardiac troponin I (cTnI) ELISA and cytokine array on heart protein lysates, respectively, where no significant changes were detected between groups (Figure S1D; Figure S2). Gross spleen weight, however, was higher in infected mice (83.3 ± 5.58 mg vs.133.5 ± 7.45 mg, mean ± SEM; Figure S1E). To measure humoral anti-MAdV-3 responses, we employed an ELISA to quantify anti-MAdV-3 IgG and find that while a significant response has been initiated at 7 d.p.i., this does not plateau until after 21 d.p.i. (Figure S1F). Together, these findings confirm that MAdV-3 is cardiotropic, representing a relevant model of acute viral myocarditis to examine mechanisms of arrhythmogenesis. While anti-viral immune responses have been initiated at 7 d.p.i, inflammatory cytokines are not detectable in the heart at the protein level. With no significant myocardial tissue damage or adaptive cellular infiltration, we conclude this stage of cardiac infection to be prior to appreciable direct host immune-mediated damage or cardiomyopathy, and faithful to reported presentation of such cardiac infections in humans^8^.

**Figure 1.**
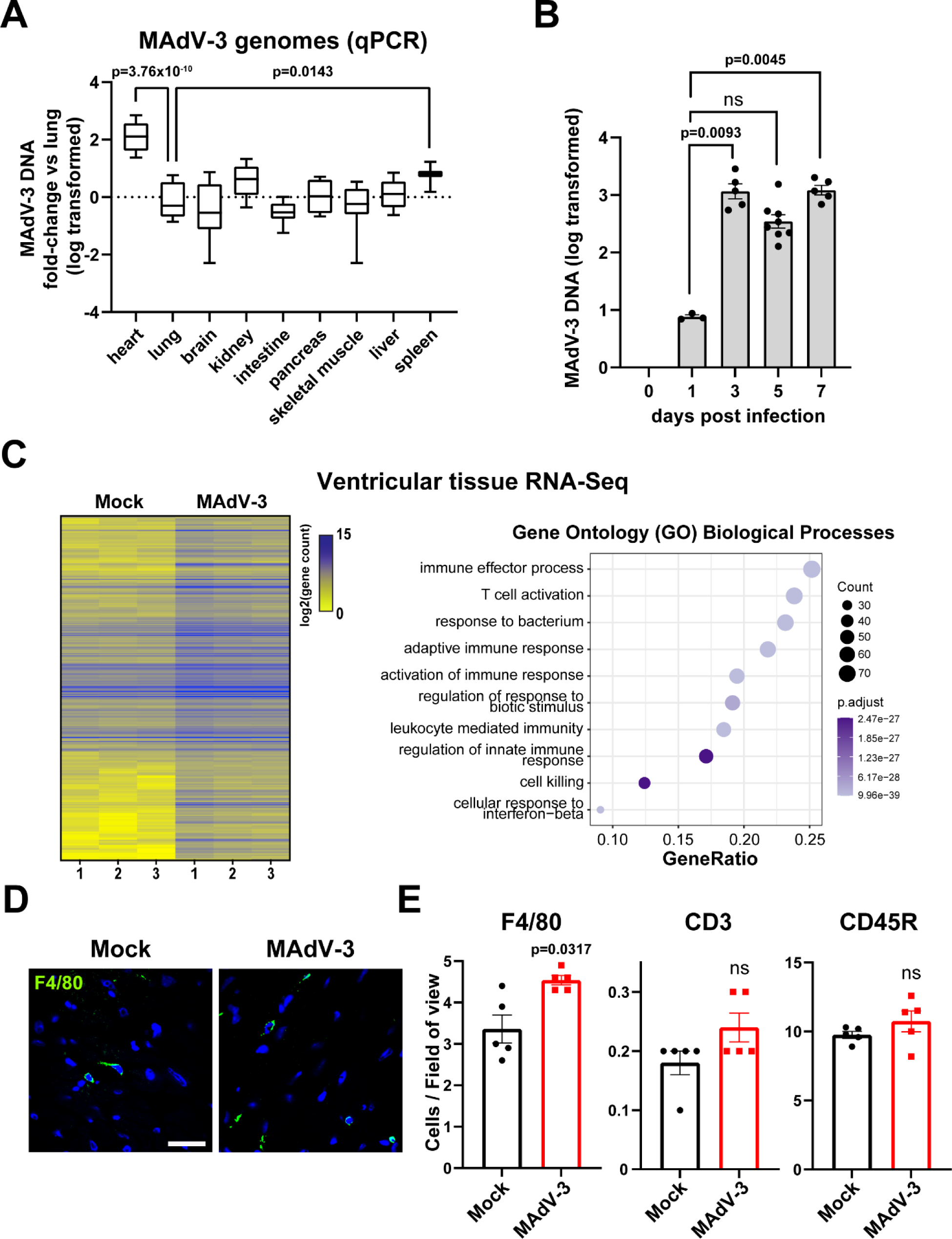
Mouse adenovirus type-3 is cardiotropic and does not invoke an appreciable infiltrative cardiac immune response during acute infection. Mice were infected retro-orbitally with 5×10^5^ i.u. MAdV-3 or saline (mock) and tissues were harvested 7 d.p.i. **A)** Using quantitative PCR, MAdV-3 viral genomes were measured across major organs and compared to organ lung tissue as a reference. (heart n=10, lung n=8, brain n=10, kidney n=10, intestine n=10, pancreas n=4, skeletal muscle n=8, liver n=12, spleen n=10). **B)** In heart tissue alone, MAdV-3 viral genomes were quantified by qPCR over a 7 day time course. **C)** RNA-Seq from ventricular tissue from control and infected animals (n=3). Left: heat map of the log2-transformed gene counts for all significant differentially expressed genes (DEGs) in infected animals compared to control (adjusted p-value < 0.05, absolute log2 fold change > 1.0). Right: gene ontology (GO) plot demonstrating the top 10 significant GO terms for biological processes identified through over-representation analysis using significant DEGs. **D)** Confocal immunofluorescence microscopy of ventricular cryosections labeled for infiltrating macrophages (F4/80, green; scale bar: 25 µm). **E)** Quantification of numbers of F4/80+, CD3+, and CD45R+ cells per field of view from confocal immunofluorescence microscopy. (n=5; 10 fields of view per heart). (**A**), One-way ANOVA with Dunnett’s multiple comparisons test ****p = 3.76×10^-10^, * p = 1.43×10^-2^ (**B**), Kruskal-Wallis with Dunnett’s multiple comparisons test ** day 3 p= 9.32×10^-3^, ** day 7 p = 4.45×10^-3^ (**E**). Mann Whitney test F4/80+ *p = 3.17×10^-2^. Data are represented as mean ± SEM.

### Cardiac conduction velocity is slowed during acute adenoviral infection in the absence of apparent cardiomyopathy

Data in Figure 1 confirm MAdV-3 cardiotropism in mice, and identify 7 d.p.i. as a timepoint where viral replication has been occurring in the heart for several days while detectable adaptive immune infiltration and/or cardiomyopathy have not yet occurred. Consistent with this, by echocardiography we find no detectable changes in ejection fraction, left ventricle developed pressure at diastole, or fractional shortening by echocardiography in infected animals compared to controls (Figure 2A). Given cases of SCD with acute viral involvement often involve essentially normal heart structure at the gross level, we next asked how cardiac electrophysiology may be affected during MAdV-3 infection. Infected and control animals were anesthetized and, using electrode pads, baseline electrocardiograms (ECGs) revealed no alterations to the PR, QRS, or QT intervals in MAdV-3-infected mice compared to mock-infected animals (Figure 2B-E). A significant prolongation of the R-R interval (114.4 ± 0.98 ms, MAdV-3 infected mean 121.4 ± 1.77 ms, mean ± SEM), indicative of bradycardia, was detected in infected animals, however (Figure 2F). To assess alterations in intercellular coupling, we next turned to measurement of conduction velocity (CV) by optical mapping. Excised hearts from infected and control animals were Langendorff-perfused and optical mapping performed to quantify longitudinal and transverse CV (CVL and CVT, respectively) during steady-state perfusion (Figure 3). While CVL was similar in both groups, we find significant CVT slowing in MAdV-3-infected hearts in comparison to mock-infected controls (38.75 ± 2.60 vs. 29.33 ± 2.88 cm/s, mean ± SEM, respectively: Figure 3C). Such reductions in ventricular conduction velocity are known to underlie arrhythmogenesis and mechanistically are typically associated with perturbation of gap junction function and/or connexin expression^17,62,66–70^.

**Figure 2.**
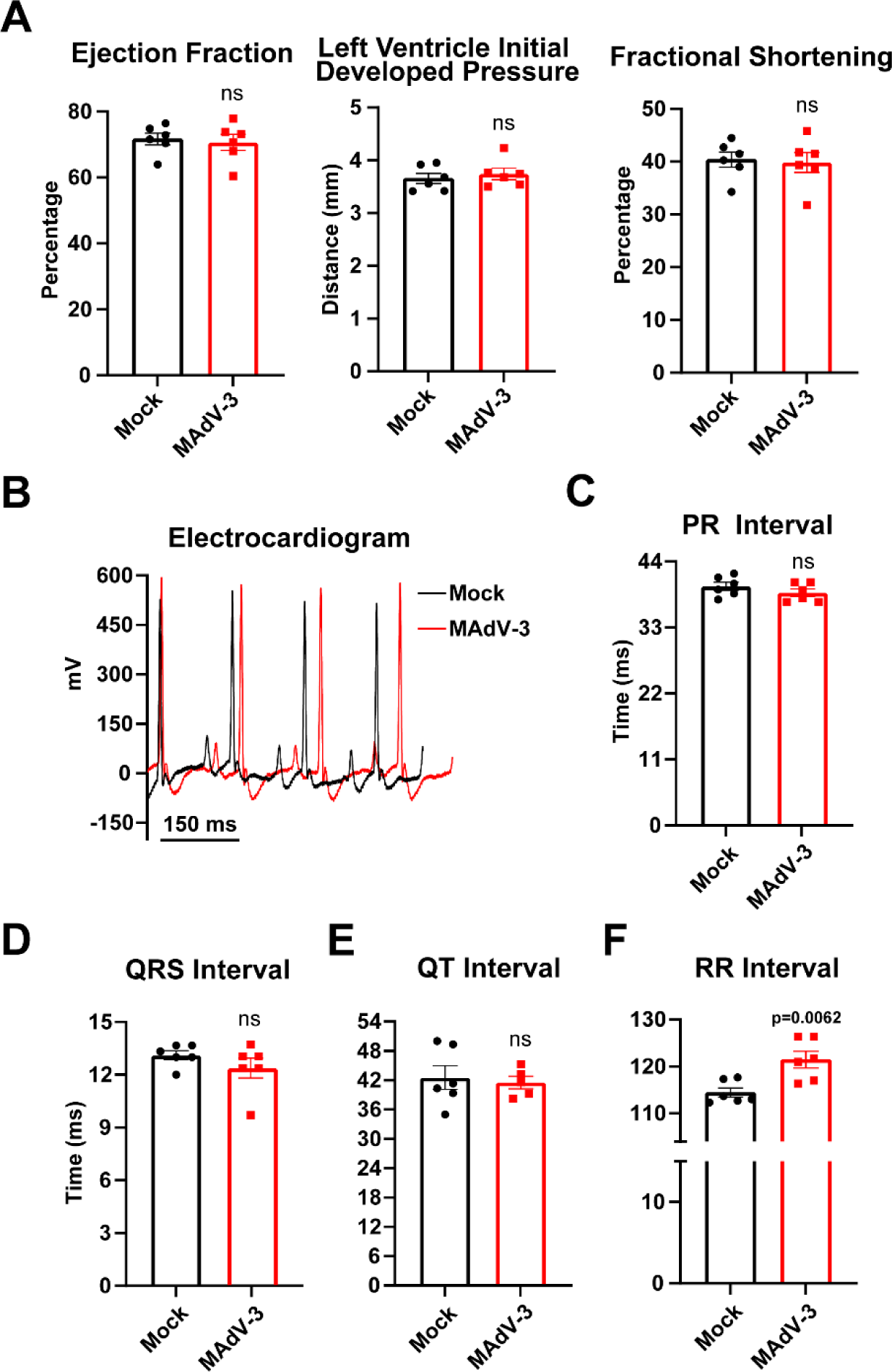
Acute cardiac adenoviral infection slows heart rate but does not alter ECG morphology or induce gross cardiomyopathy. *In vivo* echocardiography and electrocardiography were performed 7 d.p.i. in mock- or MAdV-3-infected mice. **A)** Cardiac function was measured by echocardiography on lightly anesthetized mock- and MAdV-3-infected mice (n=6). **B)** Representative *in vivo* electrocardiography measurements from lightly anesthetized mock- and MAdv-3-infected mice using paw pad electrodes. **C-F)** Quantification of PR, QRS, QT, and RR intervals from **B**. **p = 6.2×10^-3^, student’s t-test (n=6).

**Figure 3.**
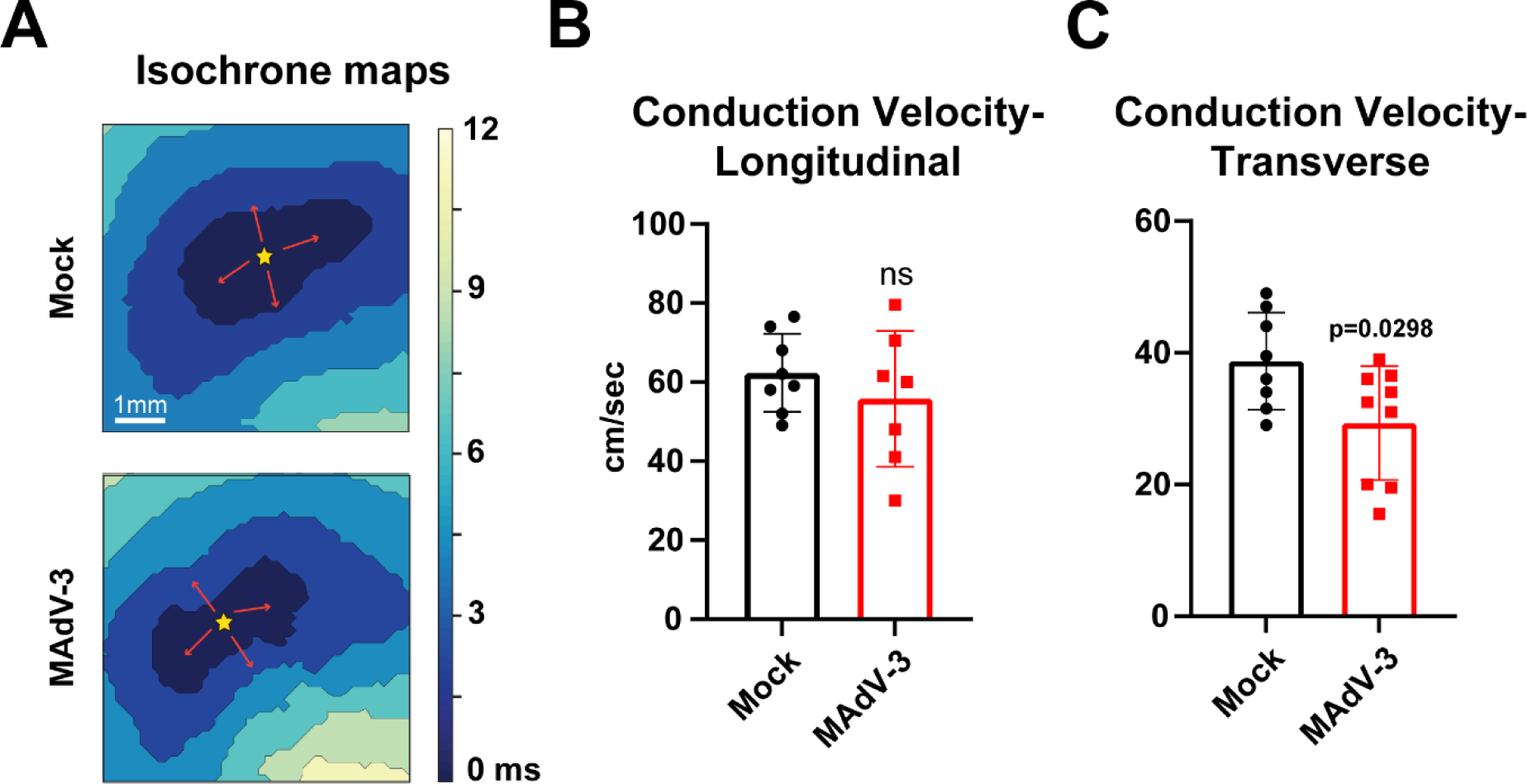
Acute cardiac adenoviral infection results in reduced ventricular conduction velocity. *Ex vivo* optical mapping was performed on Langendorff-perfused hearts 7 d.p.i. from mock- or MAdV-3-infected mice. **A)** Example isochrones maps paced at a cycle length of 150 ms with 1 ms pulse duration. **B, C)** Conduction velocity in longitudinal (CVL)and transverse (CVT) directions was quantified (n=8,9). * p = 2.98×10^-2^, student’s t-test. Data are represented as mean ± SEM.

### Aberrant connexin43 phosphorylation is induced in MAdV-3 infected hearts and primary cardiomyocytes

Having identified a conduction defect in MAdV-3 infected hearts, we next examined expression of proteins critical to cardiac electrophysiology, cardiomyocyte ID structure, and gap junction formation and maintenance. Turning to our RNA-Seq data (Figure 1C) we find no changes to expression levels of major cardiac connexins, major cardiac ion channels/subunits, or structural genes *Tjp1*, *Cdh2*, and *Pkp2* (Figure S3). We therefore investigated post-transcriptional alterations that could underlie the conduction slowing presented in Figure 3. Previously, we found a decrease of Cx43 protein localization at the cell-cell border during HAdV-5 infection in HaCaT and HiPSC-CMs^38^. To ask if gap junction localization at the ID was affected in MAdV-3 infected mouse hearts, we performed confocal immunofluorescence microscopy on myocardial cryosections to visualize Cx43 distribution in ventricular cardiomyocytes. Through quantification of Cx43 intensity at the ID, as determined through colocalization with N-cadherin^26,71,72^, we find no significant losses in gap junction localization between infected or control hearts (Figure 4A). Levels of Cx43 protein were also comparable by western blotting of lysates from MAdV-3 infected versus control cardiac tissue lysates at 7 d.p.i. (Figure 4B). Given our finding of slowed CVT without detectable alterations in Cx43 protein expression or localization, we assessed the post-translational status Cx43. Specifically, phosphorylation of Cx43^Ser368^ results in reduced channel opening probability, and our prior work with HAdV-5 identified this residue as being hyperphosphorylated during adenoviral infection of human epithelial cells^29,30,38^. Consistent with this, we find induction of Cx43^Ser368^ phosphorylation by western blotting of ventricular lysate from mice infected 7 d.p.i. with MAdV-3 in comparison to mock-infected animals (1.16 ± 0.11 vs. 1.57 ± 0.15, mean ± SEM) (Figure 4B). To ask if MAdV-3 induces Cx43^Ser368^ phosphorylation *in vitro*, we isolated primary mouse neonatal cardiomyocytes and infected them at an MOI of 10 with MadV-3 for 24 h. Induction of Cx43^Ser368^ phosphorylation was detected *in situ* by confocal immunofluorescence microscopy (Figure 4C). While control cells displayed some Cx43^368^ phosphorylation (green), most connexin detected was not phosphorylated at this residue (magenta). The majority of Cx43 detected in MAdV-3 infected cells however was positive for Ser368 phosphorylation. Quantification relative to total Cx43 reveals significant induction of Cx43^Ser368^ phosphorylation in primary neonatal cardiomyocytes compared to controls (0.98 ± 0.05 vs. 1.40 ± 0.04, mean ± SEM), importantly demonstrating infection alone is sufficient to induce this in the absence of host antiviral immune response.

**Figure 4.**
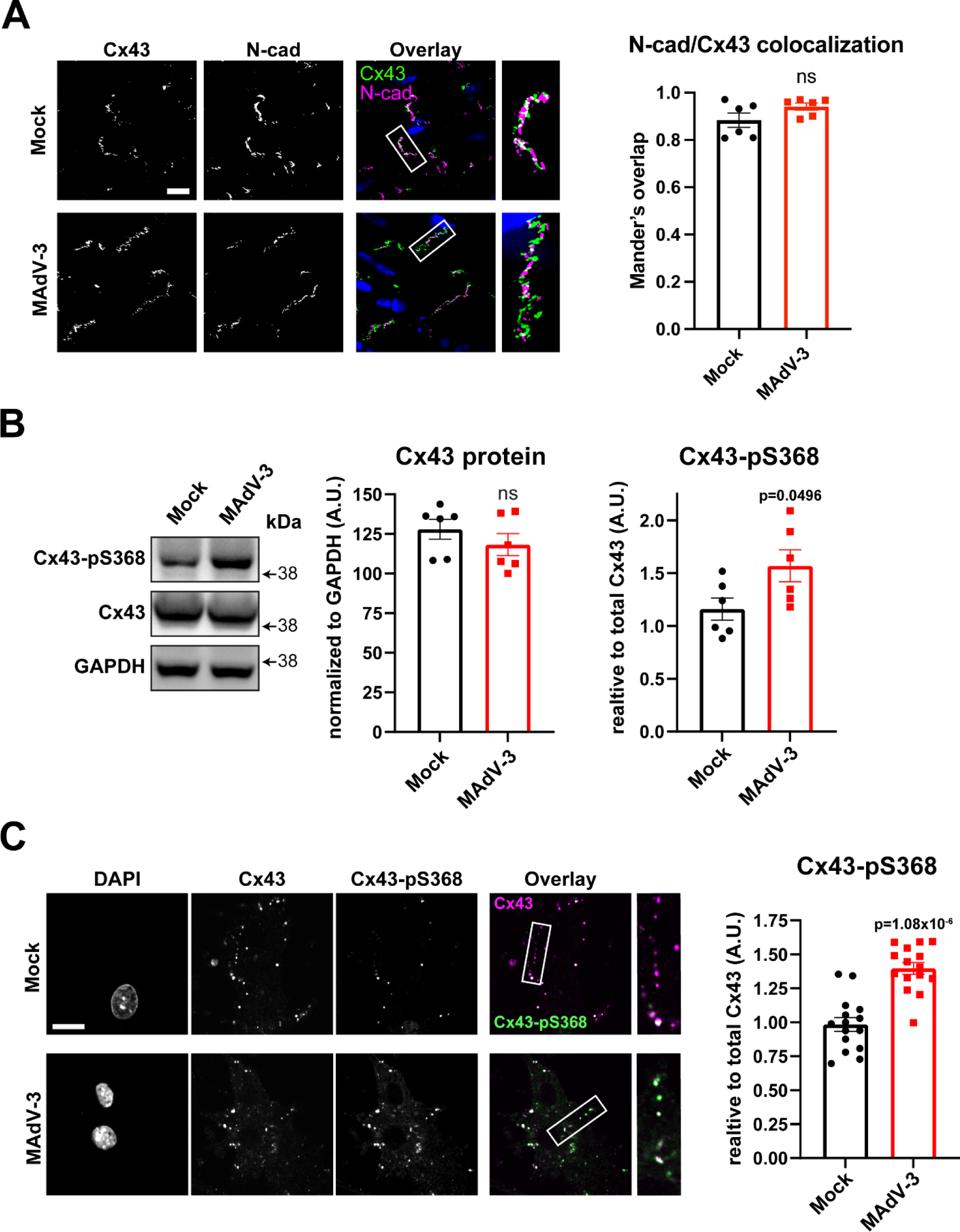
Acute adenovirus infection elicits increased connexin43 Ser368 phosphorylation in hearts and primary cardiomyocytes. Mice were infected retro-orbitally with 5×10^5^ i.u. MAdV-3 or saline (mock) and cardiac tissue was harvested 7 d.p.i. for analyses. SORA confocal microscopy of Cx43 (green) and N-cadherin (magenta) with quantification of colocalization by Mander’s coefficient (scale bar 10 µm; n=6), not significant Mann-Whitney test. Western blotting of cardiac ventricle tissue probed for phospho-Cx43 Ser368, Cx43, with GAPDH as loading control to normalize total Cx43 quantitation on left and phospho-Cx43 Ser368 normalized to total on right (n=6). student’s t-test. *p = 4.96×10^-2^, **C)** Confocal immunofluorescence microscopy of primary mouse neonatal ventricular cardiomyocytes 24 h.p.i. with MAdV-3 labeled for total Cx43 (magenta) and phospho-Cx43 Ser368 (green) (scale bar: 25 µm; n=3; **** 1.08×10^-6^ **B,C)** Data are represented as mean ± SEM.

### Adenoviral infection prolongs adult cardiomyocyte action potential duration through perturbation of potassium currents

Proper cardiac electrophysiology depends not just upon intercellular coupling and AP propagation, but the repetitive formation and resolution of consistent AP morphology through the dynamic actions of several ion channels^12^. Although no alterations in cardiac ion channel gene expression were detected by RNA-Seq and RTqPCR (Figure S3), given that we also found no change to Cx43 mRNA or protein levels, but significantly altered GJ function in infected hearts (Figure S3, Figure 4); we sought to investigate changes to electrophysiology during MAdV-3 infection. Next, we employed patch-clamping as a non-biased approach in measuring changes to cardiomyocyte electrophysiology during infection. Primary adult mouse ventricular cardiomyocytes were infected immediately post-isolation with MAdV-3 at a multiplicity of infection (MOI) of 10 or mock-infected and incubated for 24 h prior to experimentation. Importantly, infected cells were viable at this timepoint with no apparent cytopathic effect noted. Using confocal immunofluorescence microscopy, 100% infection was confirmed using MAdV-3 antiserum, where infected cells are noted to have maintained normal structure (data not shown). First, we performed whole-cell current clamp to determine if AP morphology was altered during MAdV-3 infection. While no alterations in depolarization rate (rise time/max rise slope) were detected, significant APD prolongation occurs in MAdV-3 infected cells relative to uninfected controls presented here through increased APD90 (13.87 ± 0.63 ms vs. 17.02 ± 0.67 ms, mean ± SEM) and decay time (6.03 ± 0.31 ms, 8.60 ± 0.57 ms, mean ± SEM) concomitant with decreases in max decay slope (−34.31 ± 2.82 ms vs. −26.17 ± 1.90 ms, mean ± SEM) and afterpolarization (AHP) amplitude (−1.92 ± 0.20 mV vs. −1.30 ± 0.19 mV, mean ± SEM) (Figure 5A-D). Having identified such pathological APD prolongation, we next utilized whole-cell voltage-clamp to assess individual currents and isolate the specific cardiomyocyte ion channels affected during MAdV-3 infection. No alterations in *I*_Na_ or *I*_Ca_ were detected in MAdV-3 infected ACMs (Figure 6A,B). The AP morphology changes we report occur during phases 3 and 4, however, leading us to focus in more detail on the potassium currents which primarily control proper repolarization and AP resolution. While no difference in *I*_Kr_, or *I*_Ks_ peak currents were detected between control and infected ACMs, *I*_Ks_ tail current differences (50 mV; −4.50 ± 0.64 pA/pF vs −3.15 ± 0.53 pA/pF, 55 mV; −5.03 ± 0.58 pA/pF vs. −3.16 ± 0.53 pA/pF, 60 mV; −4.03 ± 0.56 pA/pF MAdV-3 mean average −3.27 ± 0.52 pA/pF, mean difference ± SEM) and *I*_K1_ peak current (−19.80 ±1.99 pA/pF vs. −12.24 ± 1.61 pA/pF, mean ± SEM) were significantly perturbed during MAdV-3 infection (Figure 6C-E, Figure S4B,C). These data reveal potentially arrhythmogenic APD prolongation occurs in infected ventricular cardiomyocytes, compounding the electrophysiological disturbances we report affecting conduction velocity and cardiac gap junction function during acute MAdV-3 infection of the heart.

**Figure 5.**
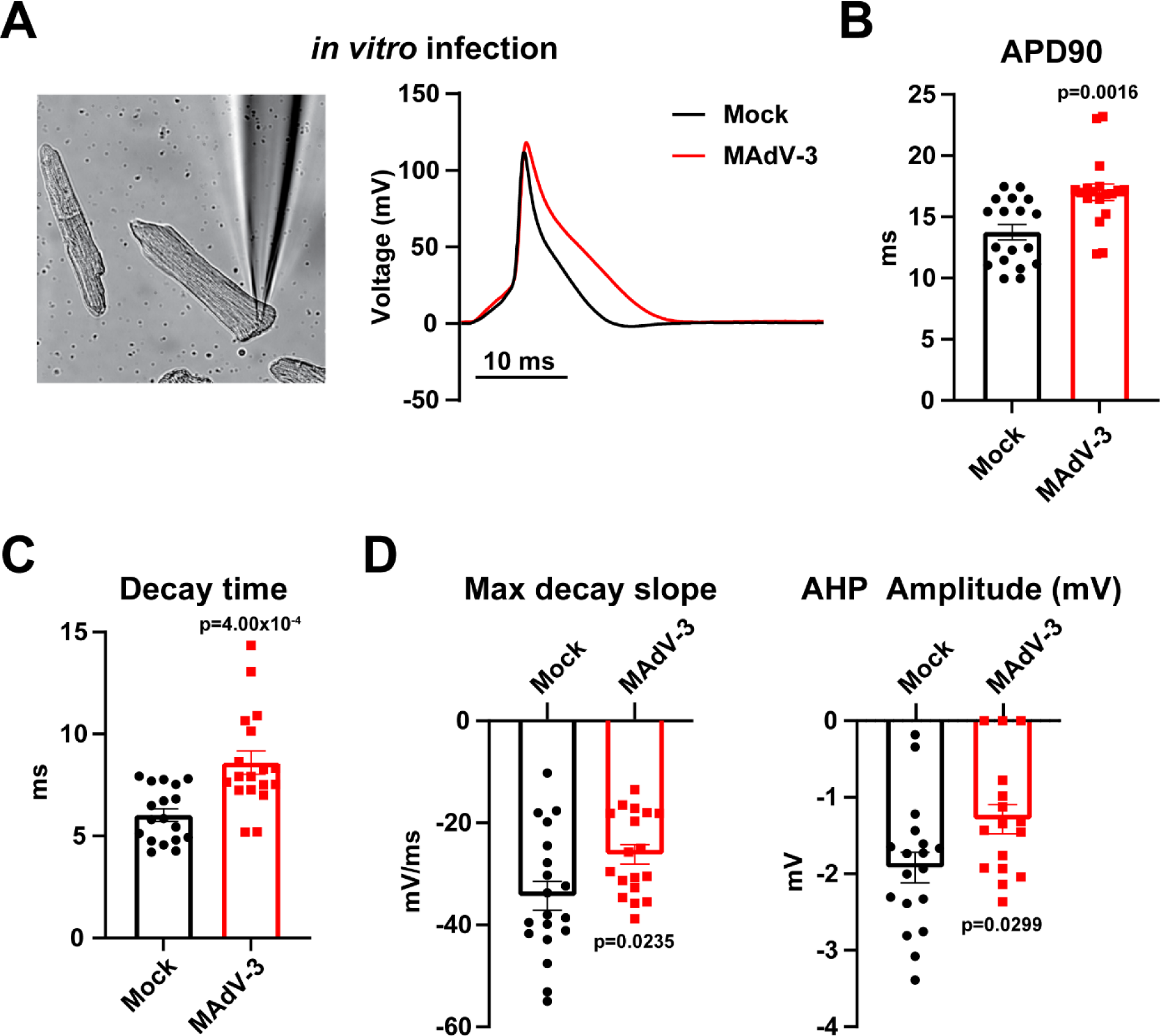
Acute mouse adenovirus infection alters ion channel gene expression *in vivo and* prolongs ventricular cardiomyocyte action potential duration *in vitro*. Isolated ACMs were either infected with MAdV-3 at an MOI 10 or not and harvested 24 h.p.i. *in vitro*. **A)** Representative image of ACMs used for patch clamp experiments with representative action potential traces recorded from uninfected (black) and MAdV-3-infected (red) cells. **B)** Action potential duration at 90% repolarization was recorded in uninfected and MAdV-3-infected ACMs (n=18). Mann-Whitney U test **p = 1.60×10^-3^ **C)** Decay time was recorded in uninfected and MAdV-3-infected ACMs (n=18). *** p = 4.00×10^-4^ **D)** Max decay slope and afterpolarization amplitude was recorded in uninfected and MAdV-3-infected ACMs (n=18). Max decay slope * p = 2.35×10^-2^, AHP amplitude * p= 2.99×10^-2^ student’s t-test. Data are represented as mean ± SEM.

**Figure 6.**
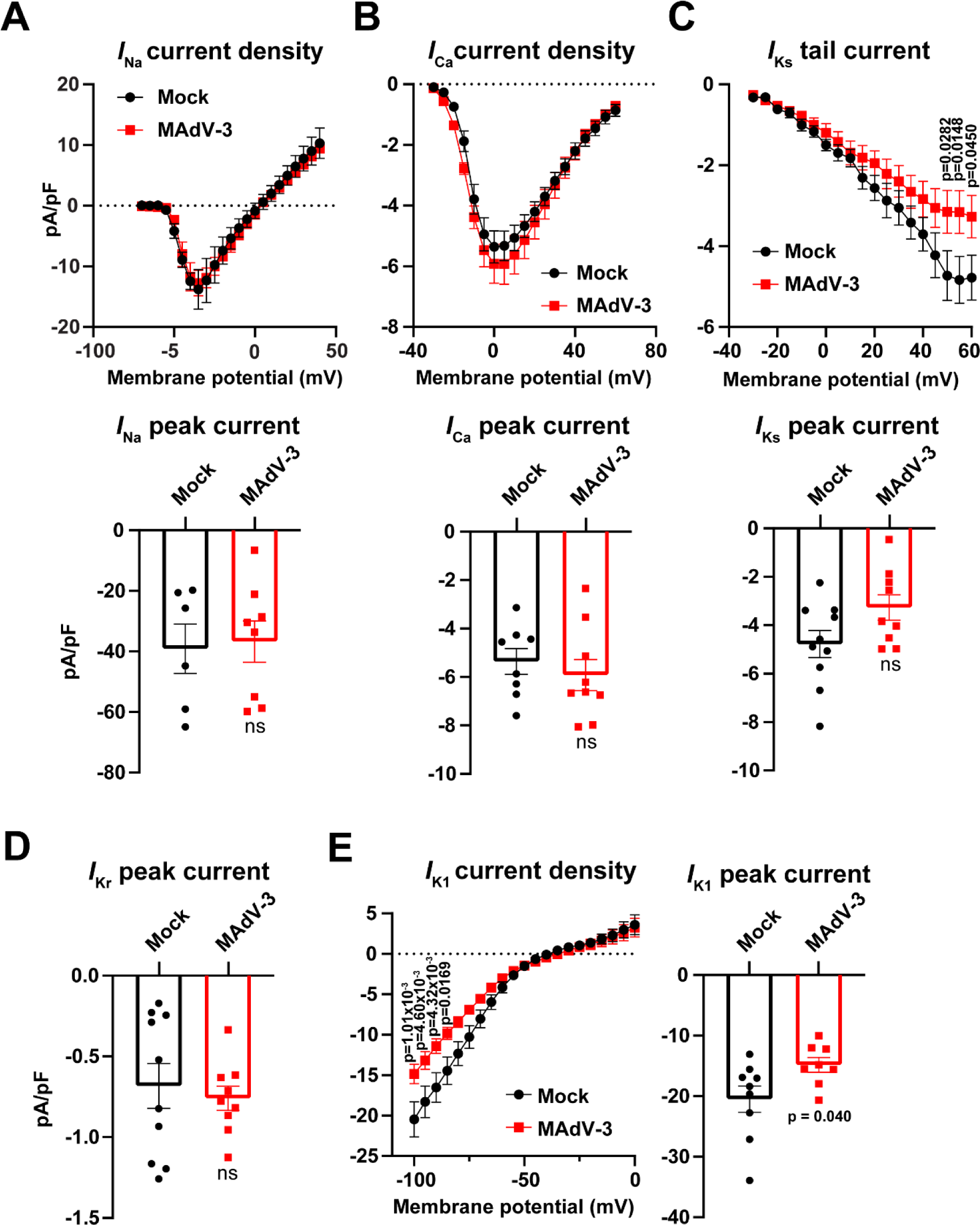
Acute mouse adenovirus infection decreases I_K1_ and I_KS_ current density in isolated adult mouse cardiomyocytes. Isolated ACMs were infected with MAdV-3 at an MOI 10 and analyzed 24 h.p.i. **A)** Current density measured isolated ACMs at 24 h.p.i. for *I*_Na_, n=6,8. **B)** Current density measured isolated ACMs at 24 h.p.i. for *I*_Ca_, n=8,9 **C)** Current density measured isolated ACMs at 24 h.p.i. for *I*_Ks_,n=10, 9, Two-way ANOVA with Šídák’s multiple comparisons test adjusted *p value= 2.82×10^-2^, 1.48×10^-2^, 4.50×10^-2^, D) Current density measured isolated ACMs at 24 h.p.i. for *I*_Kr_, n=10,9 **E**) Current density measured isolated ACMs at 24 h.p.i. for *I*_K1_, n=9,8., Two-way ANOVA with Šídák’s multiple comparisons test adjusted * p value = 1.01×10^-3^, 4.60×10^-3^, 4.32×10^-3^, 1.69×10^-2^ Peak current *I*_K1_ student’s t-test * p = 4.00×10^-2^. Data are represented as mean ±SEM.

### Human Adenovirus Type-5 increases Cx43^Ser368^ phosphorylation through PKC activation and uncouples HiPSC-derived cardiomyocytes

We previously reported that HAdV-5 disrupts expression and function of Cx43 in human epithelial cells and complexing of Cx43 with ZO-1 in HiPSC-CMs^38^. To translate our current findings of GJIC disruption and CV slowing in MAdV-3 infected mouse hearts (Figure 3, Figure 4) to a human model system, we investigated how functional intercellular coupling and Cx43 phosphorylation might be altered in HAdV-5 infected HiPSC-CMs. Matured HiPSC-CMs were cultured and infected with human adenovirus type-5 (HAdV-5) or with HAd-LacZ, a replication incompetent human adenovirus serving as control, at an MOI of 10. Phosphorylation of Cx43^Ser368^ is well established to occur via PKC^29,30,73–77^. To test if PKC activity is required for Cx43^Ser368^ phosphorylation during adenoviral infection we utilized two separate PKC inhibitors (sotrastaurin/AEB071^78,79^ ‘S’; or bisindolylmaleimide VIII/Ro 31-7549^80^ ‘B’) 1 h post infection (to ensure kinase inhibition would not hinder viral update). Just as we see with MAdV-3 in hearts and primary mouse cardiomyocytes (Figure 4B,C), HAdV-5 infection induces Cx43^Ser368^ phosphorylation at 24 h.p.i. when compared to HAd-LacZ controls (2.23 ± 0.26 vs. 12.38 ± 3.03, mean ± SEM), though not significant following the use of non-parametric testing due to small sample size. (Figure 7A). Importantly, both B and S inhibitors completely ablated this response, confirming PKC activity is necessary for Cx43^Ser368^ phosphorylation during adenoviral infection. To assess the impact of infection on electrical coupling, monolayers were loaded with the calcium sensitive Fluo4-AM 24 h.p.i. and live-cell confocal microscopy performed at 166 Hz to visualize spontaneous calcium transients. After imaging, cells were fixed and 100% infection confirmed through immunofluorescence labeling of the early HAdV-5 protein E1A (Figure 7B). While HAd-LacZ (control) HiPSC-CMs consistently displayed synchronous firing across the monolayer, uncoupling of HAdV-5 infected cells is demonstrated through significant cell-to-cell variability in time to max peak intensity (56.69 ± 8.68 ms vs. 123.8 ± 20.21 ms, mean ± SEM) (Figure 7C,D; Video S1 and Video S2). Therefore, just as we find in our MAdV-3 *in vivo* and *in vitro* studies, HAdV-5 infection disrupts cardiomyocyte electrical coupling independent of a host immune response, confirming that acute adenoviral infection generates dangerous electrical disturbances in still-viable cardiomyocytes which likely contribute to the arrhythmias of SCD attributed to acute viral myocarditis.

**Figure 7.**
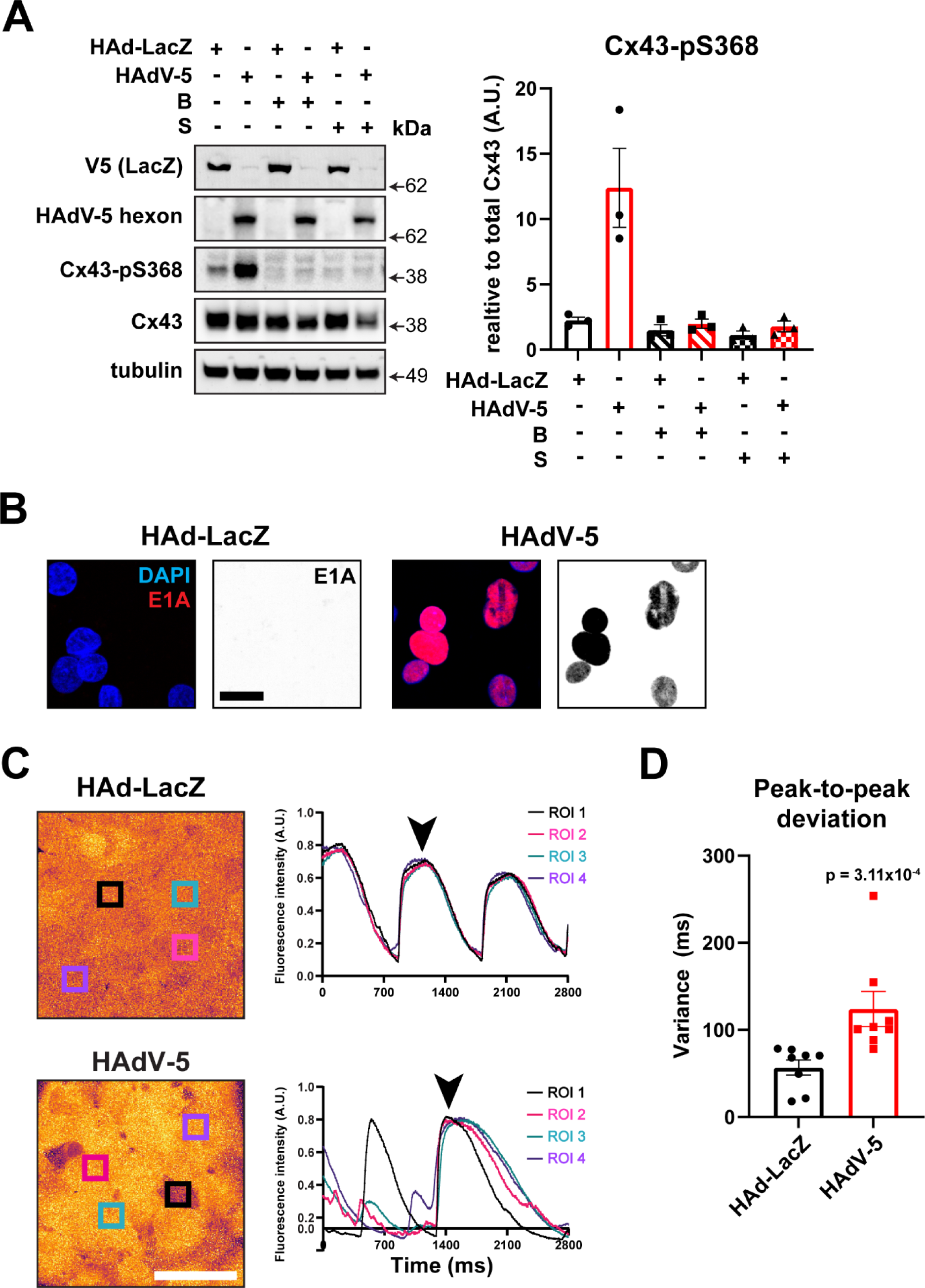
Human Adenovirus Type-5 infection increases Cx43-Ser368 phosphorylation through PKC activation and uncouples HiPSC-derived cardiomyocytes. HiPSC-CMs were infected with HAdV-LacZ or HAdV-5 at an MOI 10 and studies performed 24 h.p.i. **A)** PKC inhibitors bisindolylmaleimide VIII “B” or sotrastaurin “S” were added 1 h post infection. Western blotting of cells probed for phospho-Cx43 Ser368 with quantification normalized to total Cx43 on right (n=3), Kruskal-Wallis with Dunnett’s multiple comparisons test. **B)** Confocal immunofluorescence microscopy from fixed HiPSC-CMs to detect nuclei (blue) and Human Adenoviral protein E1A (red) (scale bar: 25 µm). **C)** Live-cell confocal microscopy 24 h.p.i. using Fluo-4 AM to measure Ca^2+^ transients and quantification of fluorescence signaling over time in one field of view. Black arrowheads indicate peak fluorescence timepoints where representative images are taken from (scale bar: 80 µm). **D)** Average peak-to-peak fluorescence intensity variance measured per cell over time (n=8,9). Mann Whitney test *** p = 3.11×10^-4^ (**D**). Data are represented as mean ± SEM.

### The phospho-null Cx43^S368A^ mutation protects against cardiac conduction slowing during acute MAdV-3 infection

We identified increases in Cx43^Ser368^ phosphorylation in MAdV-3 infected hearts where conduction slowing was occurring (Figure 3) and in HAdV-5 infected HiPSC-CMs displaying dyssynchronous firing (Figure 7). To test if prevention of Cx43^Ser368^ phosphorylation could protect infected hearts from such perturbations, we utilized transgenic mice harboring a Serine368 to Alanine mutation Cx43^81^. Complete loss of Cx43^Ser368^ phosphorylation in hearts from MAdV-3 infected Cx43^S368A^ animals was confirmed by western blot and compared to wild-type animals as a positive control (Figure 8A). As with wild-type animals, excised hearts from infected and control animals were Langendorff-perfused and optical mapping performed to quantify CVL and CVT during steady-state perfusion. We find no significant changes in CVL or CVT between control and MAdV-3-infected hearts from Cx43^S368A^ mice, indicating this mutation is protective against virally-induced conduction slowing during acute cardiac infection (Figure 8B-D).

**Figure 8.**
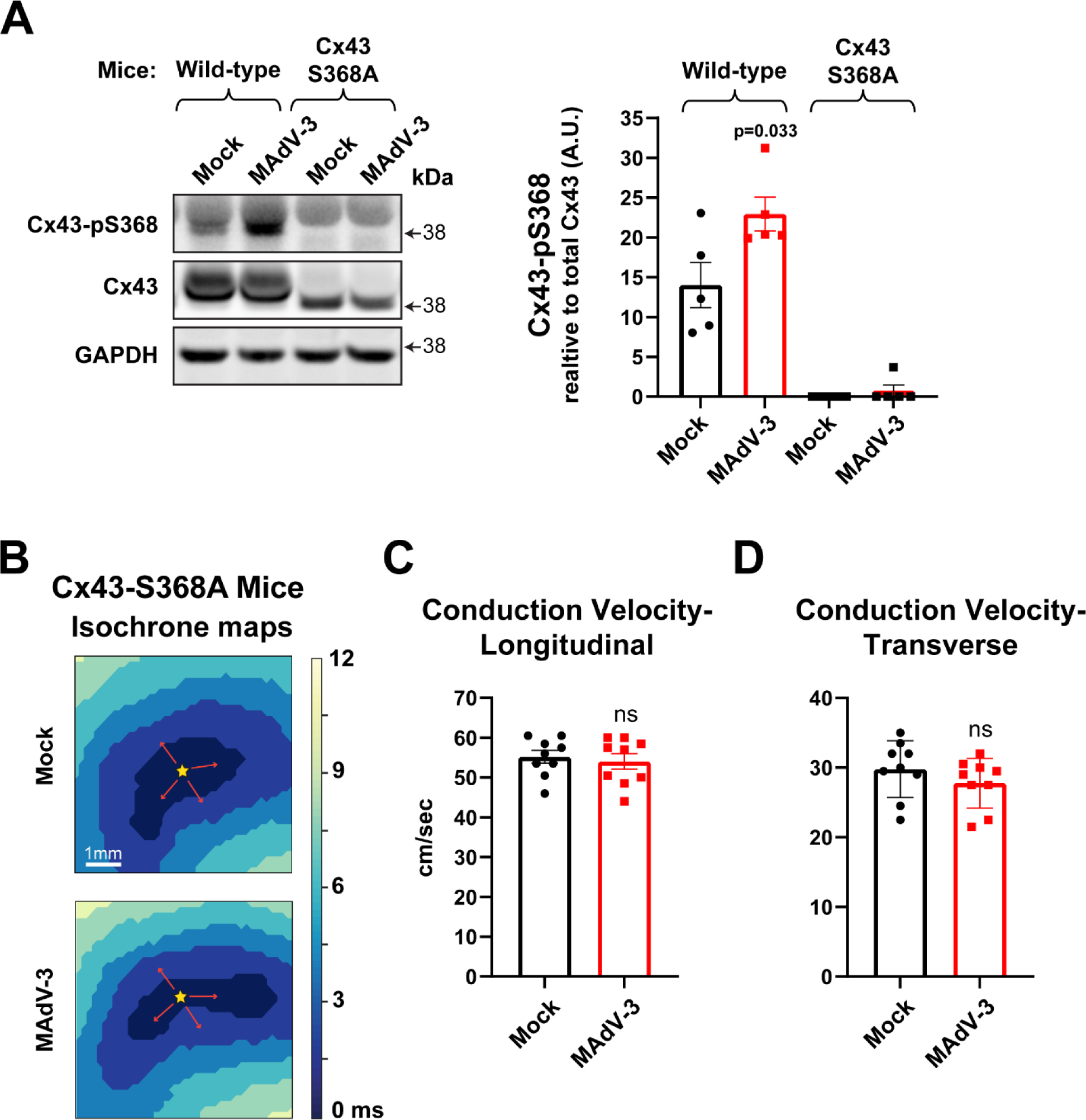
Ablation of Cx43 Ser368 phosphorylation protects against cardiac conduction slowing during acute MAdV-3 infection. Western blotting and e*x vivo* optical mapping was performed hearts 7 d.p.i. from mock- or MAdV-3-infected wild-type mice and mice harboring a Cx43-Ser368-Ala mutation. **A)** of cardiac ventricle tissue probed for phospho-Cx43 Ser368, total Cx43, and GAPDH as loading control with quantification of phospho-Cx43 Ser368 normalized to total Cx43 on right (n=5). **B)** Example isochrones maps paced at a cycle length of 150 ms with 1 ms pulse duration. **C, D)** Conduction velocity in longitudinal (CVL)and transverse (CVT) directions was quantified (n=9,9). * p = 3.30×10^-2^, (**A**), student’s t-test (**C**, **D**). Data are represented as mean ± SEM.

## Discussion

By definition, viral myocarditis encompasses inflammation and a significant contribution to either gross or molecular myocardial alterations due to infiltration of host immune cells and associated cytokine signaling^82^. There is, however, a significant gap in our understanding of pathologic events that occur during acute cardiac infection, prior to development of any detectable cardiomyopathy or mounting the infiltrative host antiviral immune response^83^. In this study, we implement MAdV-3 as a model cardiotropic adenovirus to isolate virally induced mechanisms of arrhythmogenesis during initial/acute infection that precede myocarditis or chronic disease development. While historically understudied in the context of cardiovascular disease, adenoviruses are a leading causative agent of myocarditis with the majority of cases seemingly reported in young adults and children^84^. Moreover, adenoviral genomes have been detected in cardiac tissue in cases of SCD where gross structural changes of the myocardium are not apparant^85^. While we cannot rule out immune-mediated effects completely, we report that, prior to an appreciable infiltrative adaptive immune response and/or damage to cardiac tissue; adenoviral infection decreases potassium current density leading to prolonged action potential duration with concurrent reductions in gap junction intercellular communication affecting conduction.

Connexins participate in propagation of, but are also subject to regulation by, inflammatory responses^35,36,86^. Thus, it is important in this study to parse out viral-versus host immune response-mediated induction of the arrhythmogenic substrate during acute viral infection. *In vitro*, we observe increased Cx43^Ser368^ phosphorylation in MAdV-3 infected isolated primary cardiomyocytes and HadV-5 infected HiPSC-CMs, two experimental preparations confirming phosphorylation of Cx43^Ser368^ can occur during infection independent of host immune responses (Figure 4C; Figure 7). Similarly, our patch clamping data in primary adult cardiomyocytes also demonstrate prolonged APD with disruption of potassium currents occur in infected cardiomyocytes in the absence of a host immune system (Figure 5; Figure 6). *In vivo*, we do not detect evidence of inflammation or cardiac damage, adaptive leucocyte infiltration, or increased cytokine levels at the protein level in MAdV-3 infected hearts (Figure 1D,E; Figure S1A-D; Figure S2). A primary finding of our study is hyperphosphorylation of Cx43^Ser368^ during infection to affect GJIC. Regarding specific cytokine effects, previous research has shown that increased IL-1β can prolong the QRS complex and induce Cx43^Ser368^ phosphorylation in an *ex vivo* myocarditis model in rats^87^. We find no significant difference in downstream targets of IL-1β via cytokine array 7 d.p.i. *in vivo* (Figure S2), thus we conclude this does not account for the electrical activity differences found during MAdV-3 acute infection. Additionally, VEGF signaling activates PKC and MAPK, leading to an increase in phosphorylation of Cx43^76^, however we also find no significant increase in VEGF during infection *in vivo* via cytokine array. Despite this, our RNA-Seq data from infected ventricular tissue are dominated by significantly increased mRNA signatures of the cell-intrinsic, innate, and adaptive antiviral immune responses and we report significant spleen-enlargement in infected animals (Figure 1C). Together with increased macrophage infiltration during acute MAdV-3 infection (Figure 1D,E) and a detectable humoral response (Figure S1F), we conclude that while the host antiviral response has been initiated globally and is detectable at the mRNA level in the heart, robust cardiac inflammation has not yet occurred, and the arrhythmogenic phenotypes we are observing at 7 d.p.i. herein are primarily due to active viral infection rather than immune-mediated processes.

Viruses hijack cell machinery to enable their replication while hindering host cellular and organismal immune responses^88,89^. Previous research has demonstrated that cGAS, a key signaling molecule in the innate and cell-intrinsic interferon antiviral immune response, can pass via gap junctions from infected cells to uninfected neighbors, priming and propagating the antiviral state^35^. Additionally, adaptive antiviral immune responses can be amplified in this manner by means of short linear peptide exchange via gap junctions effecting presentation of viral antigens on HLA complexes of uninfected neighboring cells that cytotoxic CD8 T-cells can engage^36^. Previously, we have reported that human adenoviral E4-ORF1 can decrease Cx43 protein expression through transcriptional repression via β-catenin, resulting in decreased gap junction intercellular communication in human epithelial cell^38^. In addition to reduced Cx43 expression, we found an increase in phosphorylation of Cx43 C-terminal sites known to be modified by AKT and PKC, retaining Cx43 at the cell membrane and reducing GJIC in cells infected with HAdV-5^38^. In this current study, we build on these findings by investigating the effects of viral modification of GJIC in the whole heart using a cardiotropic adenovirus to better understand the role of infection in precipitation of the arrhythmogenic substrate.

Consistent with our prior findings, new data from HAdV-5 infected human iPSC-CMs and MAdV-3 infected primary mouse cardiomyocytes *in vitro*, and MAdV-3 *in vivo* infections, reveal similar induction of Cx43^Ser368^ phosphorylation with concurrent reductions in gap junction function (Figure 4, Figure 7). These data demonstrate a conserved mechanism between human and mouse adenoviral infection occurring across diverse infected cellular populations (epithelial cells and cardiomyocytes) to rapidly limit intercellular communication. Perturbed gap junction coupling during adenoviral infection of the lung epithelium would presumably have limited serious pathological implications for the host. Regarding infection of the working myocardium however, such rapid disruptions in intercellular communication would likely precipitate arrhythmias of SCD during acute infection. Our data support this scenario, whereby we find Cx43^Ser368^ phosphorylation occurs in concert with both uncoupling of, and altered calcium handling in, iPSC-CMs *in vitro*, and decreased conduction velocity *in vivo* (Figure 3, Figure 7). Critically, through implementation of mice harboring Cx43^S368A^ mutation, we demonstrate that phosphorylation of Cx43^Ser368^ is necessary for conduction slowing we observe in MADV-3 infected hearts (Figure 8). Of relevance, Huke et al. previously reported a relationship between reduced intercellular coupling of cardiomyocytes, the development of focal arrhythmias, and increased Cx43 phosphorylation^90^.

Slowed conduction velocity is a well-known arrhythmogenic determinant and is closely linked to gap junction function in the heart expression^17,62,66–70^. “Previous reports have demonstrated that uncoupling of cardiomyocytes favors preferential conduction velocity decrease in the transverse direction of propagation This phenomenon is due to the higher gap junction resistance transverse to the fiber direction^91,92^. As such, the increased spatial frequency of high resistance gap junctions result in more conduction slowing transverse to fibers than along them when gap junctions are inhibited^93,94^. Thus, the reductions in conduction velocity in the transverse direction point to loss and/or alteration of gap junction function and can precipitate arrhythmias. This propensity supports our observations of a decrease of CVT in infected mice but no change in CVL as being due to gap junction dysfunction (Figure 3). We acknowledge that we observe no changes in QRS interval (Figure 2D) despite significant CVT slowing (Figure 3C). Discrepancy between ECG and optical measurements may be a result of the ECG lacking the temporal resolution to detect differences due to His-Purkinje activation of the heart in vivo, while epicardial stimulation and optical mapping slow and substantially alter activation sequences such that the same effect size differences in conduction velocity and repolarization can be detected.

Other viral agents have been shown to alter intercellular communication by way of phosphorylation of connexins during infection. Various viruses are known to decrease intercellular communication during disease, such as herpes simplex virus (HSV), feline immunodeficiency virus, and bovine papillomavirus^95–97^. Herpes simplex virus type-2 infection in rat liver epithelial cells increased serine phosphorylation 24 h.p.i. and reduce GJIC, resulting in eventual loss of Cx43 localization at the membrane^97^. In murine fibroblasts, Rous sarcoma virus, an oncogenic virus, increased phosphotyrosine and phosphoserine and reduced GJIC^98^. Human papillomavirus 16 (HPV16) protein E6 has been found to sequester Cx43 in the cytosol, limiting trafficking and gap junction formation, and potentially facilitating tumor progression. Although not directly tested, the authors report altered Cx43 phosphorylation as a likely mechanism by which HPV16 E6 achieves this^99^. Additionally, a recent study found that in a complex epithelia modeling of HPV infected reduced Cx43 protein and increased both protein expression of Cx45 and localization at the cell membrane, and this resulted in an increase of GJIC^100^. While changes to connexin phosphorylation caused by acute viral infection is most commonly studied in the context of cancer biology, clearly such changes to gap junction phosphorylation will also impact cardiomyocyte coupling across a variety of viral infections.

Adenoviral infection rapidly alters host-signaling cascades to facilitate viral invasion and replication^101,102^. Phosphorylation of Cx43^Ser368^ is well established to occur via PKC, and we confirm this in the context of adenoviral infection through implementation of two independent PKC inhibitors (Figure 7A)^29,30,73–77^. Our findings of decreased conduction velocity and increased Cx43^Ser368^ phosphorylation in the heart during adenovirus infection are in line with previous research of the interplay between viruses and PKC signaling^103,104^. For example, early in infection increased expression of adenoviral protein E1A correlates with increased PKC levels, a family of kinases whose activity is required for viral endosome escape^105,106^. Adenoviral E1A is the first protein expressed in the adenoviral life cycle and increased E1A correlates with elevated levels of viral protein along with increased PKC protein^106^. However, the E1A activation of PKC signaling in adenoviral early infection needs further delineation. PKC and downstream protein signaling regulate the cell cycle, which has been well-studied in both the fields of cancer and immune cell proliferation^107,108^. PKC exerts control over many cyclin genes, a particular note Cyclin D, via transcription and thus can uncouple cells to start cell cycle from S phase^109^. Adenoviruses push cells into S phase via E1A, E1B, and E4 protein interaction with host mechanisms, and this is required for viral replication^110–112^. Cardiomyocytes, themselves, are typically quiescent several days after birth, and therefore, are particularly recalcitrant to entering the cell cycle^113^. We must also now consider how activation of these signaling pathways shuts down gap junction intercellular communication in addition to facilitating S-phase entry. Therefore, acute adenoviral infection of the myocardium likely causes arrhythmogenesis via activation of AKT/PKC pathways which impact Cx43, and potentially other critical cardiac ion channel function.

In addition to gap junction disruption, our data provide evidence that adenoviral infection prolongs action potential duration through limiting *I*_K1_ and *I*_Ks_ in adult ventricular cardiomyocytes. Ventricular electrophysiology is therefore likely further compromised in addition to conduction delay in infected hearts (Figure 5, Figure 6). Prolonged APD can precipitate an arrhythmogenic substrate through early and/or delayed afterdepolarization in cardiomyocytes, leading to tachycardia/fibrillation^114^. The APD is determined by several distinct cardiomyocyte ionic currents such as sodium (*I*_Na_), calcium (*I*_Ca_), together with several potassium currents including *I*_K1_, *I*_Kr_, and *I*_Ks_. There is, however, a paucity of data regarding the effects of viral infection on action potential duration. Turning to models of Long QT syndromes (LQT), we can interpret how alterations to these currents contribute to sudden cardiac arrest. Anderson-Tawil syndrome (LQT7) is caused by a monogenic defect in KCNJ2 and presents as ventricular arrhythmias, prolonged QT intervals, together with physical malformations^115,116^. The ventricular arrhythmias in LQT7 are due to defects in KCNJ2, the protein responsible for the Kir2.1 inward rectifier K+ current of *I*_K1_, which lead to spontaneous action potential durations and early/delayed afterdepolarizations^116–118^. The KCNQ1 and KCNE1 proteins form the K_V_7.1 channel, creating the slow activating delayed rectifier current, *I* ^119^. Genetic modeling of LQT1 syndrome by loss-of-function in KCNQ1 causes decreased *I*_Ks_ that underlies ventricular arrhythmias and SCD^120^. Several mutations causing LQT1 impair trafficking of KCNQ1 to the cell surface, which could inform future directions in understanding the effects on *I*_Ks_ we have identified during acute adenoviral infection^121^.

Specific targeting of ion channel function has previously been described across various viral species and host cell types. In a cardiac model, CVB3 viral titers were increased in the *mayday* mutant mouse model; these mice are deficient in Kir6.1, a subunit of KATP channels, and had decreased survivability after infection^122,123^. Turning to non-cardiac excitable tissues, patch clamping experiments of neuroblastoma cells that were infected with rabies virus displayed decreased *I*_Kir_ and *I*_Na_ amplitude 3 d.p.i with no observable morphological changes^124^. While many of these studies lack a viral mechanistic cause of the change in K+ currents, it is plausible to consider this as a viral advantage, as viruses across different realms clearly and specifically alter such ionic currents during infection. Potassium channels are ubiquitously found across almost all cell types, including non-excitable cells. This included the epithelium, a common entry point for most viruses, and other cell types such as immune cells, such as macrophages and T-cells^125–128^. Interestingly, K+ channels in the epithelium are localized along the basolateral membrane, coincident with the Coxsackie adenovirus receptor (CAR) and integrins utilized by human adenoviruses for entry^126^.

It is conceivable that a common mechanism underlies the arrhythmogenic alterations to gap junctions and K+ currents we report. PKC is known to phosphorylate a multitude of host proteins relevant to our findings, including K_V_7.1 and Cx43^31^. As the kinase responsible for phosphorylation of Cx43^Ser368^, PKC activation is well-described in subsequent reduced gap junction channel opening probability^29,30,129^. PKC signaling is also known to regulate *I*_Ks_ by promoting K_V_7.1 endocytosis through increased phosphorylation of KCNE1^31^. When broadly looking at the role of kinases during ischemic stress and heart failure, PKC signaling has been found to be sufficient in *I*_K1_ reduction^130^. The commonality between negative regulation of Cx43 and *I*_K1_ and *I*_Ks_ currents by PKC strongly implicates it as a potential central kinase activated during viral infection that concurrently, and dangerously, affects distinct arms of cardiac electrophysiology. Given that PKC isoforms are known to regulate many endosomal functions, such as endocytosis and trafficking from the membrane, it is unsurprising that many viruses target this pathway for host cell invasion^131^. The direct interaction with PKC family members is vital for influenza assembly, replication machinery function, and endosomal exit^132,133^. For example, West Nile virus replication is inhibited when PKC activity is blocked and viral fusion of Respiratory Syncytial Virus (RSV) requires PKCα membrane translocation^103,134^. Thus, modification of the PKC family is a common mechanistic target across viral families.

The question remains as to how and why a DNA tumor virus such as adenovirus would infect terminally differentiated muscle, such as the working myocardium. As mentioned above, adenoviruses induce cell cycle entry to facilitate their own genome replication by arresting infected cells in S-phase^46^. Given their recalcitrance to cell cycle entry, compared to epithelial cells, it is tempting to speculate that adult cardiomyocytes are not the intended host cell for adenovirus, but when infected, precipitate an unfortunate and potentially deadly pathological process. Human adenoviruses are understood to primarily utilize CAR for binding and host cell entry, just like CVB3, although CAR-independent infection is also reported^61,135–137^. A component of the intercellular junction, CAR is expressed in several tissue types, including the heart, where it is primarily expressed on endothelial, epithelial cells, and at the cardiomyocyte ID^138,139^. In cardiomyocytes, CAR appears to play an integral role in coupling cells together for electromechanical and metabolic signaling^140,141^. The enrichment of CAR in the heart may, in part, contribute to adenoviral infection and colonization of this tissue should viremia occur, for example.

Aside from the significant body of viral myocarditis research employing CVB3, reovirus has also been developed as a model which, while typically not causing disease in humans, elicits myocarditis in neonatal mice^142^. Acute reovirus infection causes cytopathic effect specifically in cardiomyocytes, as compared to fibroblasts or epithelial cells^143^. This increases inflammation and infiltration of host immune cells, which is a common pathogenesis to that reported in CVB3 infection^144^. Mouse adenovirus Type-1 (MAdV-1) has previously been employed to investigate adenoviral-specific cardiac infection. The Weinberg group has shown significant inflammation of cardiac tissue and immune cell infiltration during chronic MAdV-1 infection^56,57^. During MAdV-1 infection, the progression of myocardial inflammation *in vivo* was assessed and viral loads in the heart peak between 7-10 d.p.i., but consistent with our findings at 7 d.p.i. with MAdV-3 (Figure 1E), significant inflammation by CD3+ cell count and cardiac troponin I serum levels were not detected until 10 d.p.i.^56^. This MAdV-1 response occurred along with high levels of IFN-γ, and the immunoproteasome induced in a dose-dependent and inverse age-dependent manner that followed IFN-γ levels^56,57,145^. These models are beneficial to understand the role of direct and indirect immune-mediated responses in the heart during infection with a virus that also afflicts the respiratory system^146^. Together, while this research has contributed much to our understanding of viral-mediated inflammation of the heart, previous studies demonstrate that there are pathogen-specific effects to the host cell at the molecular level, especially relevant to acute stages of myocarditis^83^.

In order to consistently induce MAdV-3 cardiac infection through induction of a reproducible viremia or controlled inoculum we employed the retro-orbital route of administration. While this differs from natural mechanisms preceding adenoviral myocarditis (such as infection of the gut or lung for example, prior to viremia and subsequent viremia and cardiac infection) our goals were to focus solely on pathological mechanisms in cardiac tissue once infected. Introduction of a consistent inoculum in this manner was necessary, ensuring similar exposure of cardiac muscle to virus while avoiding variability of viremia induction through other routes of administration. Having established that direct viral infection can precipitate dangerous electrophysiological remodeling in the heart, our findings provide groundwork for future studies investigating mechanisms of adenoviral cardiotropism, including why certain groups may be susceptible.

As acute infections are hypothesized to be a significant cause of sudden cardiac arrest, we sought to employ MAdV-3 as a novel acute adenoviral myocarditis model. Through studying the infected heart at early stages of viral colonization and replication, we have revealed previously underappreciated effects of viral-mediated changes to cardiomyocytes precipitating electrophysiological defects and leading to an arrhythmogenic substrate. Overall, there has been a paucity of research into direct adenoviral mechanisms that induce arrhythmogenesis and our data introduce MAdV-3 infection as a viable model with which to study adenoviral cardiac infection and myocarditis progression in the adult mouse. Relevance to human disease is confirmed through complementary work in HiPSC-CMs and HAdV-5 where we find infected cardiomyocytes undergo arrhythmogenic alterations to intercellular coupling and electrophysiology while still viable, and in the absence of an inflammatory or adaptive host immune system. Clinical relevance is also confirmed through our *in vivo* studies where, just as in human patients, we uncover an arrhythmogenic substrate in infected hearts which are free of any remarkable cardiomyopathy or inflammation^8^. Truly parsing out viral effects independent of the host inflammatory response is not possible, however, and contributions of inflammatory cytokines to our *in vivo* observations must be considered. Implementation of models to limit/modulate host immune responses may be somewhat informative here but given that in such immunocompromised contexts, viral replication and spread would be artifactually augmented, and physiologically relevance compromised. Overall, our data provide insight not just into mechanisms of arrhythmogenesis during acute cardiac adenoviral infection, but also the state of key players in cardiac electrophysiology in phases of disease preceding gross remodeling of myocarditis and progression to heart failure. In addition to informing therapeutic interventions, the ability to model acute cardiac infection will facilitate development of diagnostic approaches to identify at-risk individuals with limited clinical presentation by current screening/imaging methods.

## Acknowledgments

The authors thank Dr. Allison N. Tegge (Department of Statistics, Virginia Tech) for assisting in statistical analyses and design and Dr. Sarah Barrett (Virginia-Maryland College of Veterinary Medicine) for undertaking histopathology. The authors also thank Eric Sapp, Alex Ix, and Kenneth Young II for technical support. This work was supported by NIH NHLBI R01 grants: HL132236 and HL159512 to J.W.S.; HL138003 and HL141855 to S.P., NINDS R01 grant NS105804 and R21 grant NS128635 to S.A.S., NHLBI R35 grant HL161237P to R.G.G., NHLBI F31 grants: HL152649 to R.L.P.; HL160172 to G.A.B.; HL147438 to D.R.K., American Heart Association predoctoral fellowship award 23PRE1025476 to K.E.S., FBRI Seale Innovation Award to J.W.S., and a FBRI Lyerly Postdoctoral Excellence Award to C.M.P..

## Author contributions

J.W.S. and R.L.P. designed this study. R.L.P. performed cell and animal infections, viral propagation, primary cell isolations, RT-qPCR, qPCR, confocal immunofluorescence microscopy, western blotting, troponin ELISA, live cell imagining, echocardiography experiments and analysis. M.J.Z. analyzed RNA-Seq data, performed RT-qPCR, qPCR, confocal, western blotting, differentiated and cultured HiPSC-CMs, and assisted in all data analyses and statistics. G.A.B. and X.W. performed the optical mapping and analysis. M.T.T. performed and analyzed the echocardiograms. M.D.N. and K.E.S. performed *in vivo* infections, cryosectioning, and confocal immunofluorescence microscopy with analyses. C.M.P. performed RNA-Seq analysis and confocal immunofluorescence microscopy with analyses. S.L. performed humoral immune response ELISA as well as biochemistry and western blotting. R.G.G. provided input on optical mapping experimental design and manuscript feedback. D.R.K. and G.S.H. assisted in design and analysis of optical mapping and electrophysiology experiments. S.A.S. performed and analyzed the patch clamping experiments. S.P. provided optical mapping and electrophysiology experimental design, data analysis, and manuscript feedback. J.W.S. supervised all work and performed the cytokine array, primary cardiomyocyte isolations, and live cell imaging analysis. J.W.S., R.L.P., and M.J.Z. analyzed the data and interpretation. R.L.P. wrote the manuscript with J.W.S. with input from M.J.Z.

## Supplemental Materials

Expanded Materials and Methods

Online Figures S1-S4 and Legends

Online Videos S1 and S2

Major Resources Table

References 62-79

## Supplemental Materials

Acute adenoviral infection elicits an arrhythmogenic substrate prior to myocarditis. Rachel L. Padget, Michael J. Zeitz, Grace A. Blair, Xiaobo Wu, Michael D. North, Mira T.

Tanenbaum, Kari E. Stanley, Chelsea M. Phillips, D. Ryan King, Samy Lamouille, Robert G. Gourdie, Gregory S. Hoeker, Sharon A. Swanger, Steven Poelzing, James W. Smyth

## Expanded Materials and Methods

### Animals

All animal husbandry and experimental protocols described utilized 8-12 week old C57BL/6J male and female mice (initially obtained from The Jackson Laboratory and subsequently bred in house), were carried out under NIH guidelines, and approved by the Virginia Tech Institutional Animal Care and Use Committee. Cx43-S368A (also termed ‘PKC’ mice in prior literature) mice were a generous gift from Dr. Paul Lampe, Fred Hutchinson Cancer Center, Seattle, WA^62^.

### Cell culture

Normal murine mammary gland (NMuMG), A549, (ATCC), 3T6, 293A, and 293FT (Thermo Fisher) were maintained in Dulbecco’s Modified Eagle Medium, high glucose with L-glutamine and sodium pyruvate (Genesee Scientific) supplemented with 10% FBS, non-essential amino acids (Gibco) and Mycozap Plus-CL (Lonza). Primary mouse neonatal ventricular cardiomyocytes were isolated as previously described^63^ and maintained in F12/DMEM 50/50 supplemented with 2 % FBS (Gibco), insulin–transferrin–sodium selenite media supplement, 10 μM BrdU, 20 μM cytosine β-darabinofuranoside (Sigma-Aldrich), and Mycozap-PR (Lonza). BrdU and cytosine β-darabinofuranoside were removed 24 h prior to MAdV-3 infection studies.

### HiPSC-CM differentiation, expansion, and maturation

HiPSCs (Cell Applications) were differentiated into cardiomyocytes as previously described^64^, with modifications. Briefly, iPSCs were seeded in E8 medium (Thermo Fisher) into matrigel (Corning) coated (1:99) 12-well plates with daily media changes for 3 days. Following this, on day 0, medium was changed to CDM3, as defined in^65^ with 3 µM CHIR-99021 (Selleckchem) in RPMI for 2 days. On day 2, medium was changed to CDM3 with 2 µM Wnt-C59 (Cayman). On day 4, medium was replaced with CDM3. On day 6, medium was changed to RPMI1640 + B27 (Thermo Fisher) (-insulin) and refreshed on day 8. On day 10, medium was changed to RPMI1640 without glucose + B27. On day 12, cardiomyocytes were expanded as in^66^. Cells were lifted with TrypLE Select Enzyme 10X (Thermo Fisher), washed with PBS 20% FBS and centrifuged at 200 x *g* for 4 min, resuspended in RPMI 1640 + B27 1X with 10% Knock Out Serum Replacement (Gibco) and Thiazovivin 1.0 μM (Selleckchem), and transferred into 10 cm dishes. 24 h later medium was replaced with RPMI 1640 + B27 1X supplemented with 2 μM CHIR-99021 (Selleckchem). Cells were further expanded for 4 passages prior to cryopreservation. IPSC-CMs were verified by the presence of cardiac troponin-T, N-cadherin, and spontaneous contraction under phase microscopy. Prior to experiments, IPSC-CMs were matured in a defined maturation medium composed of oxidative substrates and low glucose for 3 weeks with media changes every 3 days, as previously reported^67^.

### Virus propagation and titering

MAdV-3 (a generous gift from D. H. Krüger, M.D., Ph.D., Institute of Med. Virology, University Hospital Charité, Berlin) was propagated in NMuMG cells. MAdV-3 was purified by PEGylation and CsCl ultracentrifugation as previously described, and titer was determined by immunofluorescence confocal microscopy in 3T6 cells (Thermo Fisher)^38,68^. HAd5 (ATCC) was propagated in A549 cells and AdlacZ was generated according to manufacturer’s instructions from pAd/CMV/V5-GW/LacZ (Thermo Fisher) for purification by CsCl ultracentrifugation and titering in 293A cells as previously described^38,68^.

### Adult mouse cardiomyocyte isolations

Adult mouse cardiomyocytes (ACMs) were isolated as previously described^69^. In brief, mice were anesthetized with isoflurane and administered a 500 U heparin i.p. injection (Sigma Aldrich). Hearts were excised and cannulated within 4 min post excision on a Langendorff apparatus utilizing pH 7.4 perfusion buffer for 3 min comprising (in mM, all from Sigma Aldrich): NaCl (120.4), KCl (12.7), KH_2_PO_4_ (0.6), Na_2_HPO_4_ (0.6), MgSO_4_-7H_2_O (1.2), NaHCO_3_ (4.6), Taurine (30), Glucose (5.5), 2,3-Butanedione 2-monoxime (BDM, 10), and HEPES (10). Perfusate was then switched to perfusion buffer including 2.4 mg/ml Type II Collagenase (Worthington Biochemical) followed with 3 µM CaCl_2_ reintroduction when 20 mL of solution remain to perfuse. Aorta, vena cava, and atria were removed, and the ventricles were mechanical broken apart into single cells. Cells were centrifuged at 30 x *g* for 3 min, and resuspended in three sequential stop buffers comprised of perfusion buffer and FBS + 10, 40, 100 µM CaCl_2_ reintroduction steps were completed and cells were plated on dishes coated with 100 µg/ml laminin (Gibco) in cardiac myocyte media (CMM, ScienCell) with provided cardiac myocyte growth supplement, 5% FBS, 50 mM BDM, and MyocoZap Plus-PR (Lonza), which was replaced on cardiac myocytes at 2 h after plating.

### Infections and inhibitor treatments

#### In vivo

8-12 week old C57BL/6 male and female mice were anesthetized and inoculated via retro-orbital injection with 5×10^5^ infectious units (i.u.) of MAdV-3 diluted in 0.9 % sterile saline (Teknova) or saline solution alone for control animals, using U-100 Insulin syringes (BD). Mice were monitored daily for weight loss and assessment of fur ruffling/neurological symptoms, none of which were noted in the work described herein.

#### In vitro

Infections were performed at a multiplicity of infection (M.O.I.) of 10 i.u./cell as previously described^38^. For PKC-inhibition experiments IPSC-CMs were seeded in 12-well culture plates at 90 % confluency and matured. Cells were treated with 5 µM Bisindolylmaleimide VIII (E2839; Selleckchem) or 5 µM sotrastaurin (S2791; Selleckchem) 1 h post infection with Ad5 or LacZ. After 24 h protein was harvested, and western blotting performed as described below.

### RNA-Seq

Ventricles were separated from atria and snap frozen for RNA-seq processing by GeneWiz, Inc. who performed alignment and initial analyses. Sequencing was performed by HiSeq 2×150 PE HO HiSeq 2×150 bp configuration. DESeq2 was used to compare gene expression between groups of samples. The Wald test was used to generate p-values and log2 fold changes. Genes with an adjusted p-value < 0.05 and absolute log2 fold change > 1 were called as differentially expressed. Log2 transformed normalized gene counts of significant differentially expressed genes were hierarchically clustered using the heatmap program clustergrammer^70^, with the Scipy library in Python, using default settings of cosine distance and average linkage. Over-representation analysis was conducted with significant differentially expressed genes using the R/Bioconductor package clusterProfiler^71^, with p-values adjusted using the Benjamini-Hochberg method. The top ten most significant gene ontology terms for biological processes were plotted in order of descending gene ratio.

### Cytokine array

Mouse hearts were excised, rinsed in ice-cold PBS and snap frozen for downstream use in cytokine array. Ventricle tissue was lysed and array performed according to manufacturer’s instructions (ab133993; Abcam) and developed and quantified utilizing a Chemidoc MP imaging system with Image Lab software (Bio-Rad).

### Cardiac Troponin I and humoral immune response ELISAs

Mouse serum was processed from blood collected from the mandibular vein and heart excision of mock and MAdV-3 infected animals. Cardiac Troponin I (cTNI) serum levels were measured using a sandwich ELISA according to manufacturer’s instructions (ab246529, Abcam). The antiviral humoral response was detected as previously described with modifications utilizing the OptEIA Reagent Set B (BD Biosciences)^72^. Briefly, 96-well plates were coated with UV-inactivated purified MAdV-3 at 1 × 10^4^ i.u./well overnight in 100 mM sodium carbonate coating buffer at 4 °C overnight an washed three times before blocking for 1 h in assay diluent. Serum samples were diluted 1/500 in assay diluent and 100 µl added to each well in triplicate, and incubated for 1 h at room temperature before 3 washes. Secondary antibody (HRP conjugated goat anti-mouse IgG, 1/3,000; Abcam) was diluted in assay diluent, 100 µl added to each well, and incubated for 1 at room temperature before 3 washes and substrate development according to manufacturer’s instructions. Data for both ELISAs were acquired utilizing a SpectraMax i3 plate reader (Molecular Devices).

### Echocardiography

Echocardiography was performed and analyzed using VisualSonics Vevo 660 High-Resolution Imaging System with a MS250 transducer at 7 days post-infection. Mice were anesthetized with 2 % isoflurane in 2 % supplemental O_2_ with long axis and M-mode images recorded, and surface electrodes measured cardiac electrical activity as previously reported^73^. Heart rate, electrocardiography, and left ventricle (LV) dimensions of ejection fractions (EF %), fractional shortening (FS %), left ventricular developed pressure (LVDP) at diastole were measured.

### Histology

Histological preparation, blood counts, and serum chemistry analysis were performed according to standard operating procedures in the Virginia Tech Animal Laboratory Services (ViTALs) diagnostic laboratory accredited by the American Association of Veterinary Laboratory Diagnosticians (AAVLD) at Virginia-Maryland College of Veterinary Medicine. Tissues for histology were fixed in 10 % neutral buffered formalin (VWR) for 24 h, then trimmed, processed, paraffin-embedded, and sectioned at 5 µm prior to staining with hematoxylin and eosin (Abcam). Histopathologic examination was performed in a blinded fashion by a board-certified veterinary pathologist. Tissues evaluated included heart, brain, lungs, thymus, salivary glands, esophagus, tongue, eyes, spleen, adipose tissue, liver, kidneys, adrenal glands, pancreas, stomach, duodenum, jejunum, ileum, colon, skeletal muscle, bone, bone marrow, and haired skin.

### Tissue DNA extraction and qPCR for viral genomes

For viral tropism, tissue was homogenized in TRIzol (Thermo Fisher) and DNA was extracted according to manufacturer’s instructions in combination with ethanol precipitation. For viral time course experiment, hearts were homogenized on a Bead Mill 4 (Thermo Fisher) and digested using DNeasy blood and tissue kit (Qiagen) according to manufacturer’s instructions. For detection of viral genomes, qPCR was performed with SYBR Select Master Mix for CFX (Thermo Fisher) on a QuantStudio 6 Flex System (Thermo Fisher). Cellular genome DNA reference gene C1 primer forward-5’-CCT TCC AGT TGA GTC AGT GG-3’ reverse-5’-CTG CTG CTG TTG GGT ACT TC-3’. For detection of MAdV-3 viral genomes, MAdV3 qPCR-2.FORWARD: GTG GGA GGA TTT GAG GAG ATG; MAdV3 qPCR-2.REVERSE: ATG TAG CAC AGG CAT TAG TAG G were employed.

### RNA extraction and RT-qPCR

Tissue was homogenized in TRIzol (Thermo Fisher) and clarified by phenol-chloroform phase separation according to manufacturer’s instructions. RNA was further purified with the PureLink RNA Mini Kit (Thermo Fisher) and DNA was removed by on column DNase digestion. cDNA was generated with iScript Reverse Transcription Supermix for RT-qPCR (Bio-Rad) according to manufacturer’s instructions. Real-time PCR was performed with SYBR Select Master Mix for CFX (Thermo Fisher) on a QuantStudio 6 Flex System employing the following primers: *Hprt* Mm.PT.39a.22214828 IDT; *Cacna1c*, Mm.PT.58.8608981 IDT; *Scn5a*, Mm.PT.58.30418114 IDT; *Kcnh2*, Mm.PT.58.42865919 IDT; *Cdh2*, Mm.PT.58.12378183 IDT; *Pkp2*, Mm.PT.58.11914062 IDT; T*jp1*, Mm.PT.58.12952721 IDT; *Gja1*, Mm.PT.58.5955325 IDT; *Gjc1*, Mm.PT.58.8383900 IDT.

### Langendorff heart preparations and optical mapping

Animals were anesthetized and cervical dislocation performed upon loss of peripheral stimuli response and immediately followed by thoracotomy and cardiac excision. Hearts were cannulated and Langendorff-perfused within 4 min, as previously described^74^. Hearts were perfused with a pH = 7.4 baseline crystalloid solution containing (in mM): NaCl (144.5), CaCl_2_ (1.8), NaOH (5.5), KCl (4), Dextrose (5.5), MgCl_2_ (1), HEPES (10), NaH_2_PO_4_ (1.2). Solutions were bubbled with 100% O2 for the duration of the experiment and run at a constant flow to maintain a pressure of ∼70-90 mmHg, with perfusates and the tissue bath maintained at 37 °C. Following a 15 min stabilization period, hearts were perfused with 8 µM Di-4-ANEPPS (Biotium), excess dye was washed out over a 20 min period, and images acquired during subsequent perfusion. The electromechanical uncoupler blebbistatin (10 µM, ApexBio Tech), as well as slight pressure on the posterior surface of the heart, were used to minimize cardiac motion and stabilize the heart against the glass of the imaging window. Hearts were paced with a unipolar silver wire placed on the anterior epicardium and a reference wire placed in the back of the bath. Stimulation at 1.5 X the diastolic threshold was applied at a basic cycle length of 150 ms and 1-2 ms pulse duration. Hearts were paced for 30 sec before each image was collected. Action potentials were recorded as previously described^75^. Briefly, the anterior epicardium of the heart was illuminated by a light-emitting diode (LED) light source (LEX3G, SciMedia) with a 520/35 nm band-pass filter (Brightline), and reflected onto the heart via epi-illumination using a 565 nm dichroic mirror (Chroma Technology). Emitted light was collected via a tandem lens system and transmitted through a 610 nm long-pass filter (Andover Corp.) before detection with a MiCAM Ultima L-type CMOS or 02CMOS camera (SciMedia). Action potentials were recorded at a sample rate of 1 kHz for a duration of 2 sec during intrinsic activity and steady state pacing. Conduction velocity (CV) was collected in the transverse and longitudinal direction (CVT and CVL, respectively), and was calculated as previously described^74^. Briefly, activation time for each pixel was designated as the maximum rate of rise of the optical action potential. Conduction in each direction was quantified by selecting vectors up to 5 pixels away and within an angle of ± 8° from a user-defined line indicating the direction of action potential propagation. Conduction vectors within 2 rows of the pacing site were excluded to reduce pacing artifacts. Data are presented as representative isochrone maps with 2 ms time steps and summary data (mean ± SEM).

### Western blotting

Mouse hearts were homogenized in RIPA buffer (50 mM Tris pH 7.4, 150 mM NaCl, 1 mM EDTA, 1 % Triton X-100, 1 % sodium deoxycholate, 2 mM NaF, 200 μM Na_3_VO_4_, 0.1 % sodium dodecyl sulfate, 5 mM *_N_*-ethylmaleimide; all from Sigma Aldrich) supplemented with HALT protease and phosphatase inhibitor cocktail (Thermo Fisher). Lysates were clarified by sonication and centrifugation and concentration determined by DC protein assay (Bio-Rad). 4X Bolt LDS sample buffer (Thermo Fisher) supplemented with 400 mM DTT (Sigma Aldrich) was added to samples then heated to 70 °C for 10 min and subjected to SDS-PAGE using NuPAGE Bis-Tris 4-12 % gradient gels with MES (Thermo Fisher) running buffer according to manufacturer’s instructions. Proteins were transferred to LF-PVDF (Bio-Rad) membrane and fixed in methanol and dried. PVDF membranes were reactivated in methanol followed by blocking in 5 % nonfat milk (Carnation) or 5 % bovine serum albumin (Fisher Scientific) in TNT buffer (0.1 % Tween 20, 150 mM NaCl, 50 mM Tris pH 8.0) for 1 h at room temperature.

Primary antibodies were incubated overnight at 4 °C utilizing: rabbit anti-Cx43 (1:3,000; Sigma-Aldrich), mouse anti-GAPDH (1:2,000; Santa Cruz Biotechnology), mouse anti-α-tubulin (1:3,000; Sigma Aldrich), rabbit mAb [D3H8Q] anti-V5 (1:1,000; Cell Signaling Technology), rabbit anti-HAdV-5 (1:5,000; Abcam), and rabbit mAb [D6W8P] anti-phospho-Cx43^Ser368^ (1:1,000; Cell Signaling Technology). Membranes were washed six times with 1 X TNT before secondary antibody labeling. Membranes were labeled for 1 h at room temperature with secondary antibodies conjugated with AlexaFluor555 or AlexaFluor647 (1:5,000; Thermo Fisher) or HRP (1:5,000; Abcam). For HRP labeling (phospho-Cx43^Ser368^), membranes were stripped for 30 min in ReBlot Plus Strong Antibody Stripping Solution (Sigma Aldrich) before reprobing for total Cx43. Membranes were imaged and quantified utilizing a Chemidoc MP imaging system with Image Lab software (Bio-Rad).

### Patch clamping

Electrophysiological recordings from dissociated cardiomyocytes were performed at room temperature using whole-cell voltage-clamp and current-clamp techniques. Recordings were made with an Axopatch 200B amplifier, Digidata 1550B digitizer, and pClamp 11 software (Molecular Devices) with a 20 kHz sampling rate and low-pass filtering at 5 kHz. Series resistance was monitored and compensated at > 70 % during voltage-clamp recordings. Recording electrodes with a tip resistance of 2-4 MΩ were pulled from borosilicate glass. Action potentials and potassium currents (*I*_K1_, *I*_Ks_, and *I*_Kr_) were recorded with an internal pipette solution containing (mM): K-aspartic acid (120), KCl (20), NaCl (10), MgCl_2_ (2), and HEPES (5) brought to pH 7.3 with KOH, and extracellular solution containing (mM): NaCl (120.4), KCl (12.7), KH_2_PO_4_ (0.6), Na_2_HPO_4_ (0.6), MgSO_4_-7H_2_O (1.2), NaHCO_3_ (4.6), Taurine (30), Glucose (5.5), and HEPES (10) brought to pH 7.4 with HCl. Sodium current and calcium currents were recorded with an internal pipette solution containing (mM): CsMeSO_3_(120), tetraethylammonium chloride (20), MgCl_2_ (2), HEPES (10), EGTA (10), and Mg-ATP (4) brought to pH 7.3 with CsOH. Sodium currents were recorded in extracellular solution containing (mM): NaCl (25), N-methyl-D-glucamine (120), CaCl_2_ (1.8), MgCl_2_ (1.8), glucose (10), and HEPES (10), pH 7.3, supplemented with 1 μM nisoldipine to block voltage-gated calcium channels. Calcium currents were recorded in extracellular solution containing (mM): NaCl (137), CsCl (5.4), MgCl_2_ (1.8), CaCl_2_ (2), glucose (10), and HEPES (10), pH 7.3, supplemented with 10 μM TTX to block voltage-gated sodium channels.

Resting membrane potential and action potentials were recorded in current-clamp mode. RMP was measured by averaging the membrane potential across 5 seconds with no current injection. Action potentials were recorded with the membrane potential held at −90 mV by a constant bias current and elicited by 2 ms current pulses between 200 pA to 2 nA at 1 sec intervals. Action potential shape analysis was performed on action potentials elicited in response to the lowest current injection using the Action Potential module in Clampfit 11 (Molecular Devices). *I*_K_, *I*_Na_, and *I*_Ca_ were recorded in voltage-clamp mode, and the 10 mV liquid junction potential was corrected for during the recordings. *I*_K1_ was elicited from a holding potential of −40 mV with 2.5 sec depolarizing voltage steps from −100 mV to 0 mV and quantified by measuring the peak steady-state current amplitude during the voltage steps. To measure *I*_Ks_ and *I*_Kr_, cardiomyocytes were held at a holding potential of −40 mV holding potential, given depolarizing voltage pulses from −30 to 60 mV for 2.5 sec, and then stepped back to −40 mV in the absence and presence of 5 μM E4031 (Cayman chemical). *I*_Ks_ was determined by measuring the peak amplitude of the tail current in the presence of E4031, and *I*_Kr_ was determined by measuring the peak amplitude of the E4031-sensitive current. *I*_Na_ was elicited from a holding potential of −90 mV with 20 ms depolarizing steps from −80 mV to +60 mV. *I*_Ca_ was recorded from a holding potential of −40 mV with 300 ms depolarizing steps from −30 mV to +60 mV. *I*_Na_ and *I*_Ca_ were determined by measuring the peak current amplitude relative to baseline during the voltage steps. Three runs were completed per cell for all recordings.

### Immunofluorescence microscopy

As previously described^38,76,77^, cells were fixed with −20 °C methanol for 5 min, washed, and blocked with 5 % normal goat or donkey serum (Thermo Fisher) containing 0.1 % Triton X-100 (Sigma Aldrich) which was increased to 0.5 % for ACMs for 1 h at room temperature. Primary antibody labeling was performed for 1 h at room temperature using; rabbit anti-Cx43 (1:3,000; Sigma), goat anti-Cx43 (1:250; Abcam), rabbit mAb [D6W8P] anti-phospho-Cx43^Ser368^ (1:1,000; Cell Signaling Technology), mouse anti-N-Cadherin (1:250; BD Biosciences), mouse anti-Adenovirus E1A [M73] (1:2,000; Abcam), rat anti-F4/80 (1:250; Abcam), rabbit anti-CD3 (1:250; Novus), or goat anti-CD45R (1:500; Novus). Cells was washed 6 times prior to labeling for 1 h at room temperature with fluorescently-conjugated secondary antibodies raised in goat (AlexaFluor 488, 555, or 647; 1:500; Thermo Fisher) or donkey (AlexaFluor 488 or 647; 1:500; Jackson Labs and CF568; 1:500; Biotium) and counterstained with DAPI (Thermo Fisher). Immunofluorescence of tissue was performed on 8 µm acetone-fixed cryosections from snap frozen tissue in OCT as previously described^77^. The labeling protocol was the same as for cells with the modification of overnight incubation with primary antibodies at 4 °C overnight and inclusion of 0.05% Tween 20 (Fisher Scientific) in 2 initial washes following primary and secondary antibody incubations. Confocal imaging was performed on SoRa Spinning Disk (Nikon) or Opterra swept field confocal microscope (Bruker) and image processing was undertaken using FIJI software (NIH)^78^. SoRa images were deconvolved and processed utilizing NIS-Elements Advanced Research software (Nikon) and Bruker Opterra confocal image processing was undertaken using FIJI software (NIH)^78^. Colocalization of N-cadherin and Cx43 was assessed via Pearson’s and Mander’s correlation coefficient across the Z-stack. Antibody specificity was confirmed through inclusion of secondary-only controls. Additionally, optimization and experimental cardiac CD3, CD45R, and F4/80 immunostaining was performed in tandem with spleen tissue sections to confirm both specificity and successful labeling. Acquisitions and analyses were performed blind with fields of view chosen at random within areas of interest (e.g. left ventricle) and representative images presented were chosen to reflect that concluded by averaged summary data.

### Live calcium transients imaging

HiPSC-CMs were plated into 35 mm glass bottom culture dishes (Cellvis) and maintained as described above to form monolayers and mature. Cells were infected with HAdV-5 at an MOI of 10 and loaded with Fluo-4AM (Thermo Fisher) diluted in Tyrode’s solution containing (mmol/L): NaCl (140), KCl (5.4), MgCl_2_ (1), sodium pyruvate (2), CaCl_2_ (1), glucose (10), HEPES (10), pH adjusted to 7.4 using NaOH at room temperature^79^. Live cell imaging was performed utilizing an Opterra swept field confocal microscope (Bruker) with a 20x objective with an acquisition rate of 166.67 Hz for at least 6 fields of view per well encompassing several spontaneous firings. Fluorescence from image sequences encompassing one field of view was normalized in FIJI^78^ by generating maximum (MaxIP) and minimum (MinIP) intensity projections, and subtracting MinIP from the original image sequence and MaxIP. The resulting image sequence was divided by the result of MaxIP-MinIP. Regions of interest (ROIs) were defined over individual cells and fluorescence intensity plotted over time. ROIs were selected at random from cells that were firing during a 2.916 second-long live cell imaging series. ROI were kept size-consistent across all image analysis and acquired data from the cytosolic space of cells. Standard deviation of timepoint for peak fluorescence intensity of one firing from 4 cells in each sequence was calculated as a measure of synchronicity.

### Statistics

All experiments were repeated at least three times and blinding of data was implemented prior to quantification (see figure legends for specific replicate values). All technical replicates are averaged into single data points, so only biological replicates are presented. Data are presented as mean ± SEM. Statistical analysis was conducted with GraphPad Prism (GraphPad Software, Inc.). A value of p<0.05 was considered statistically significant.

**Supplemental Figure 1.**
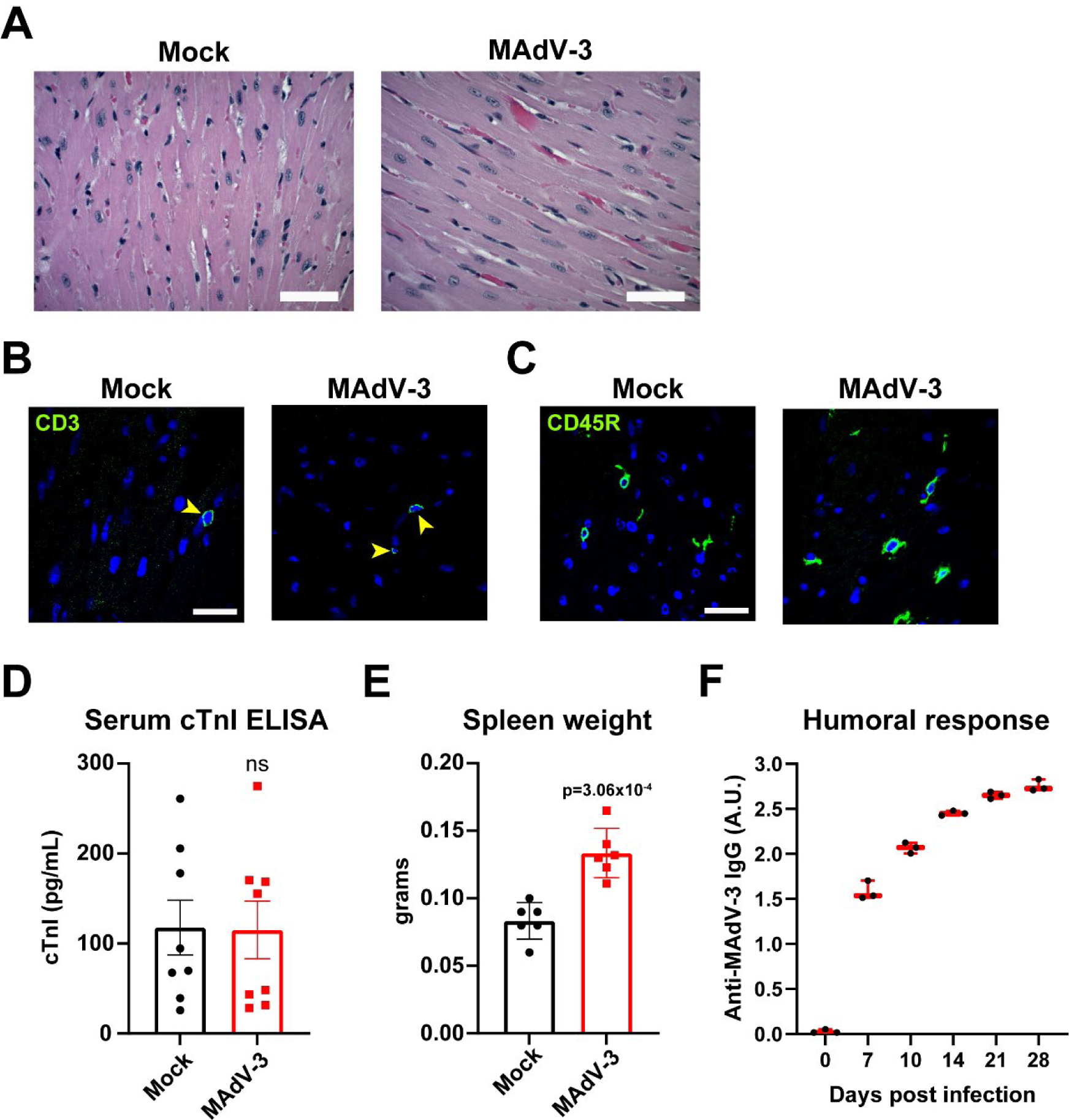
Acute MAdV-3 infection does not induce inflammatory cardiomyopathy but does increase spleen weight and elicit a humoral response. Mice were infected retro-orbitally with 5×10^5^ i.u. MAdV-3 or saline (mock) and spleen tissue or serum was harvested 7 d.p.i. for analyses. **A)** Histopathology by H&E staining on paraffin embedded cardiac sections (scale bar: 50 µm). **B)** Confocal immunofluorescence microscopy of ventricular cryosections labeled for infiltrating T-lymphocytes (CD3, green; scale bar: 25 µm). **C)** Confocal immunofluorescence microscopy of ventricular cryosections labeled for infiltrating B-lymphocytes (CD45R, green; scale bar: 25 µm). **D)** Cardiac troponin I ELISA from circulating blood serum of mock or MAdV-3 infected mice was quantified (n=8). **E)** Gross spleen weights (n=6). **F)** Anti-MAdV-3 mouse IgG ELISA from serum to detect humoral immune responses (n=3). ***p = 3.06×10^-4^, student’s t-test. Data are represented as mean ± SEM.

**Supplemental Figure 2.**
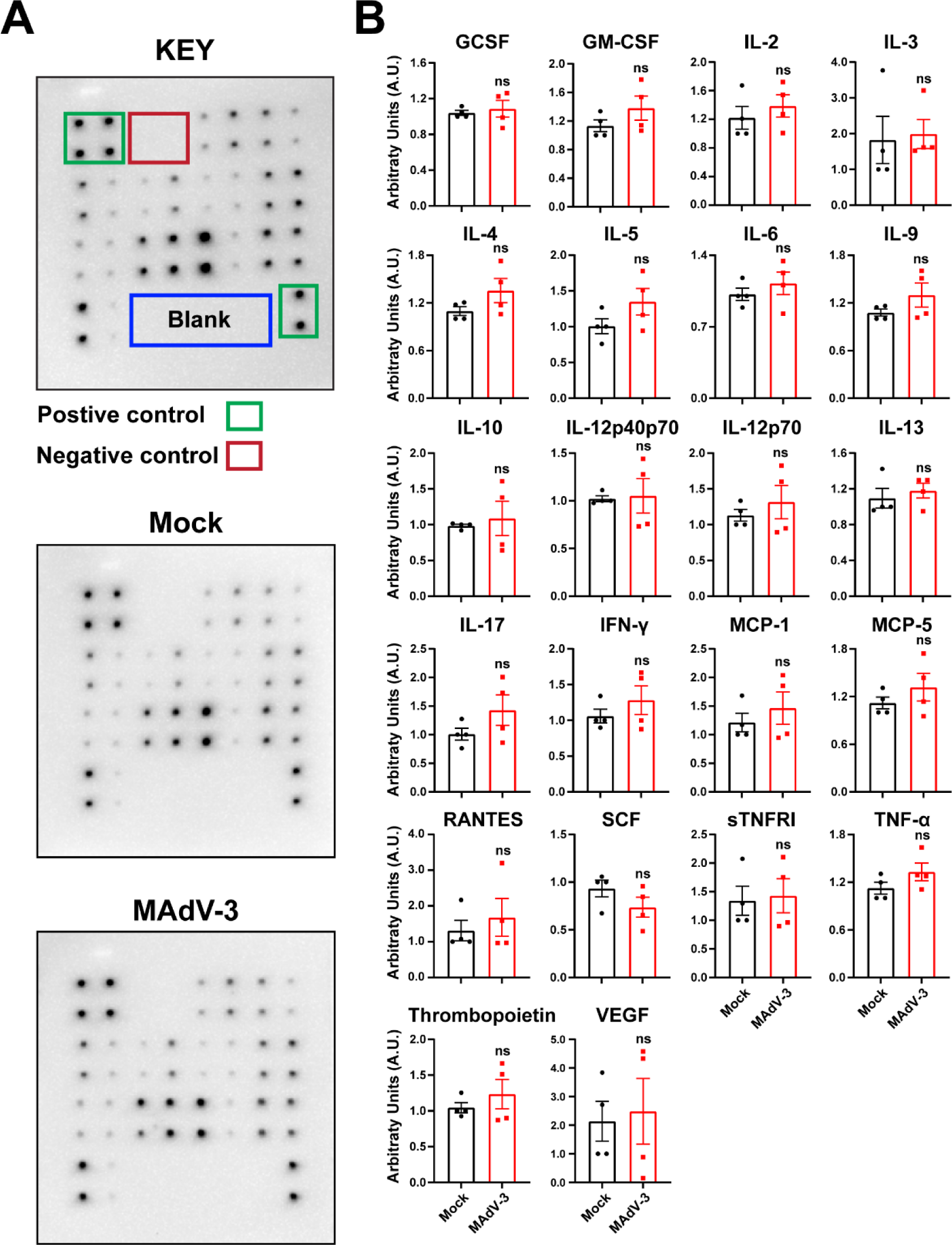
Acute MAdV-3 infection does not elicit a detectable cytokine response in the heart 7 days post infection. Mice were infected retro-orbitally with 5×10^5^ i.u. MAdV-3 or saline (mock) and cardiac tissue was harvested 7 d.p.i. for analyses. **A)** Representative whole blot images from cytokine array performed on cardiac tissue lysate. **B)** Quantification of blots from cytokine array (n=4). Student’s t-test. Data are represented as mean ± SEM.

**Supplemental Figure 3.**
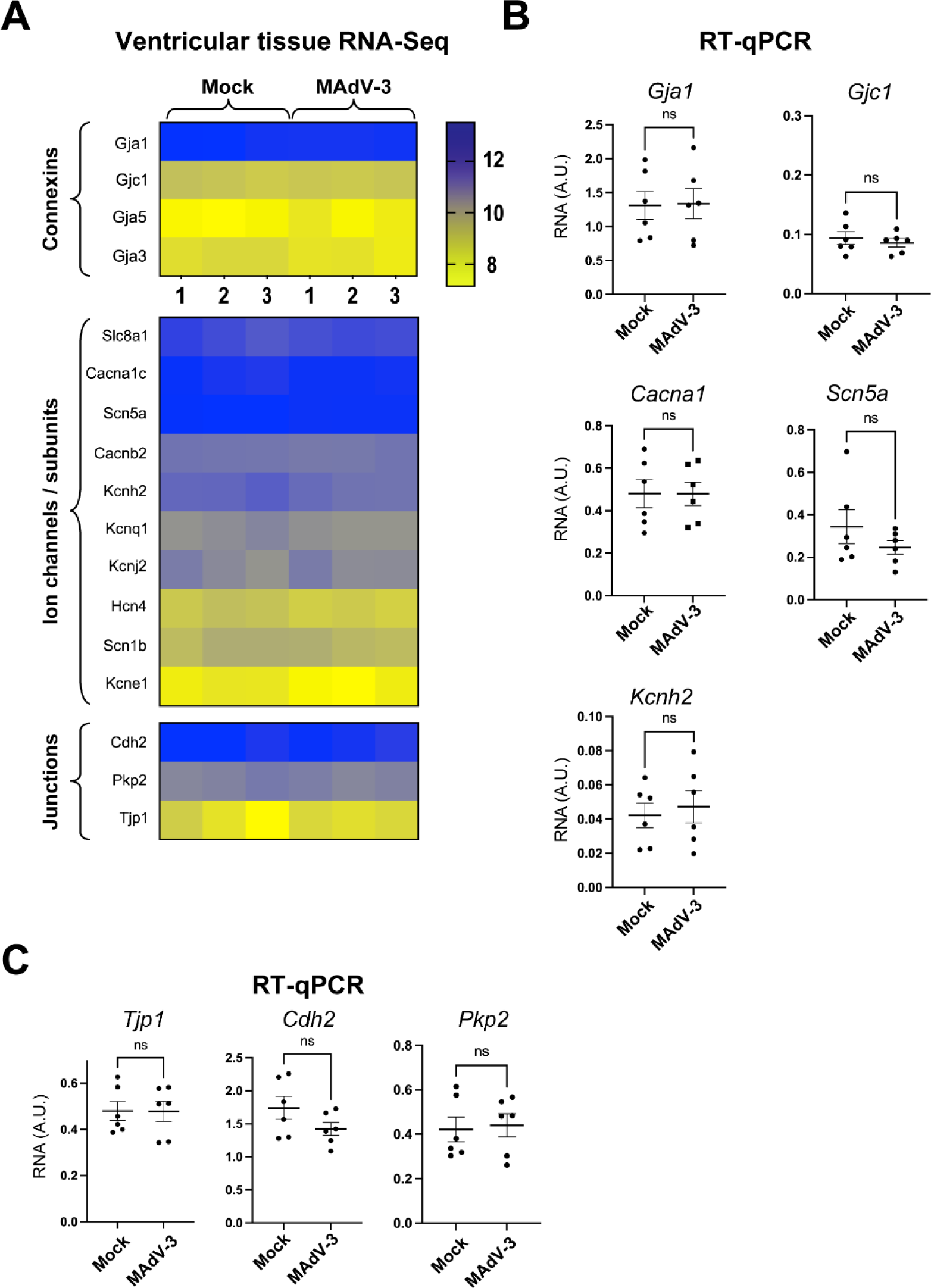
Acute adenovirus infection does not alter gene expression of major cardiac electrophysiological and structural proteins. Mice were infected retro-orbitally with 5×10^5^ i.u. MAdV-3 or saline (mock) and cardiac tissue was harvested 7 d.p.i. for analyses. **A)** Heat map of RNA-Seq data for major cardiac connexins, ion channels and subunits, and intercalated disc encoding mRNAs (n=3). **B)** RT-qPCR of heart tissue mRNA for *Gja1*, *Gjc1*, *Cacna1*, *Scn5a*, and *Kcnh2* (n=6). **C)** RT-qPCR of heart tissue mRNA for major cardiac scaffolding genes *Cdh2*, *Pkp2*, and *Tjp1*, (n=6). student’s t-test. Data are represented as mean ± SEM.

**Supplemental Figure 4.**
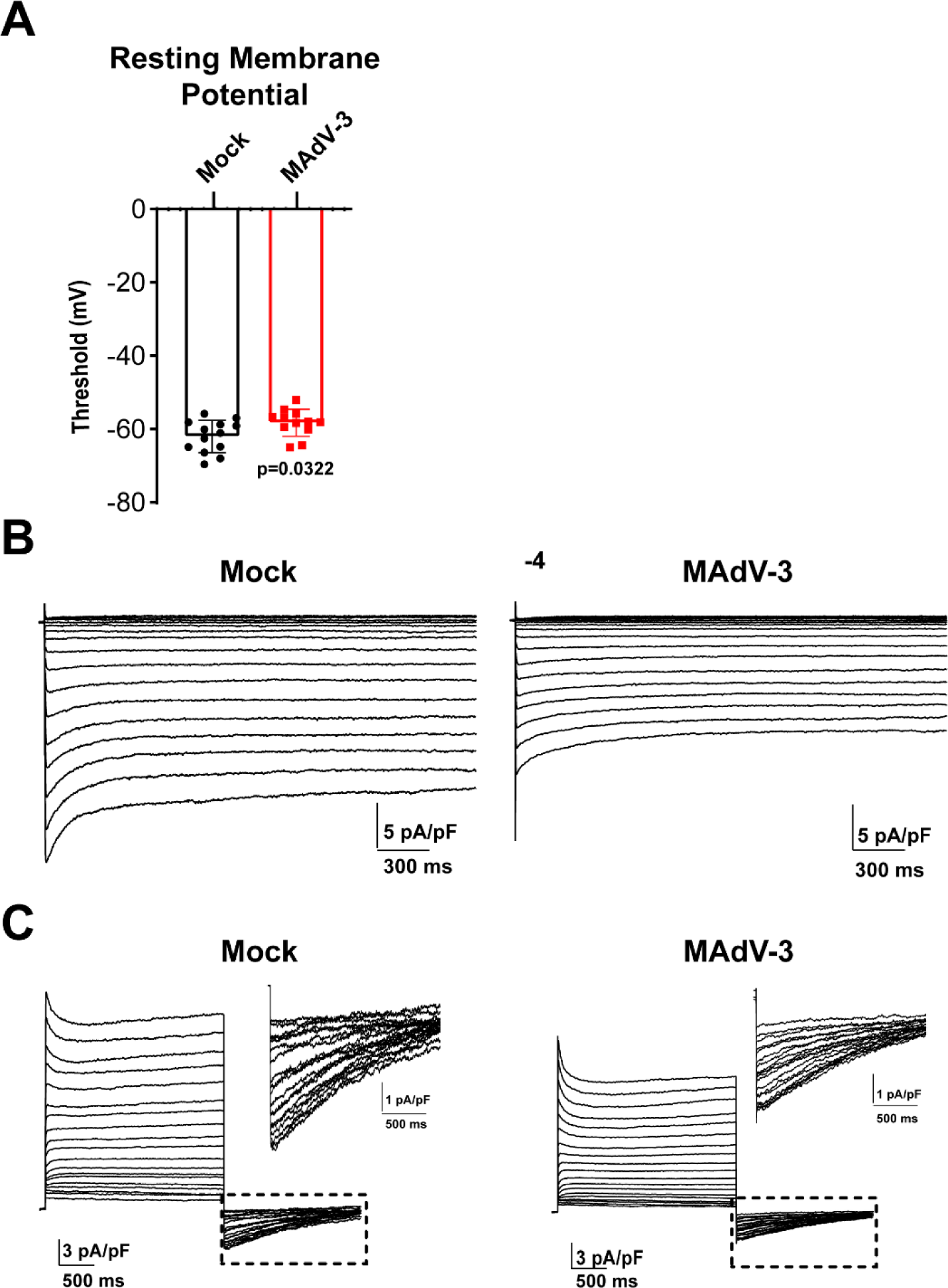
Cellular resting membrane potentials and raw traces of *I*_K1_ and *I*_Ks_ currents. Isolated ACMs were infected with MAdV-3 at MOI 10 and were patch clamped at 24 h.p.i., **A)** Resting membrane potentials recorded from mock or infected isolated ACMs (student’s t-test p = 3.22×10^-2^ n=13,12). **B)** Full current traces of *I*_K1_ in uninfected and infected cardiomyocytes. **C)** *I*_Ks_ insert currents measured in the presence of 5 µM E4031, n=19.

**Supplemental Movies 1 and 2.** Related to Figure 7. HiPSC-CMs were infected with HAdV-LacZ or HAdV-5 at an MOI 10 and harvested 24 h.p.i. Live-cell confocal microscopy 24 h.p.i. using Fluo-4 AM to measure Ca^2+^ transients was performed at 166 Hz.

## Non-standard Abbreviations and Acronyms

ACM: Adult mouse ventricular cardiomyocyte
APD: Action potential duration
cTNI: Cardiac Troponin I
Cx43: Connexin43
CV: Conduction velocity
CVB3: Coxsackievirus B3
CVL: Longitudinal conduction velocity
CVT: Transverse conduction velocity
DCM: Dilated cardiomyopathy
ECG: Electrocardiogram
EF: Ejection fraction
FS: Fractional shortening
GJIC: Gap junction intercellular communication
HPV: Human papillomavirus
HSV: Herpes simplex virus
HAdV-5: Human adenovirus type-5
HF: Heart failure
HiPSC-CM: Human induced pluripotent stem cell derived cardiomyocyte
HRP: Horseradish peroxidase
ID: Intercalated disc
IFN: Interferon
LQT: Long QT syndrome
LV: Left ventricle
LVDP: Left ventricular developed pressure
MAdV-3: Mouse adenovirus type-3
MOI: Multiplicity of infection
NMuMG: Normal murine mammary gland epithelial cells
NMVCM: Neonatal mouse ventricular cardiomyocyte
SCD: Sudden cardiac death
ZO-1: Zonula occludens-1

